# Multi-timescale motor circuit dynamics underlies adaptive and efficient exploratory behavior

**DOI:** 10.1101/2024.11.18.624064

**Authors:** Pinjie Li, Heng Zhang, Yifan Su, Jiaqi Wang, Louis Tao, Quan Wen

## Abstract

Motor systems must balance flexibility and structure to enable efficient and adaptive movements. Traditional hierarchical descending models struggle to explain the intrinsic dynamism in natural behaviors. Here, we investigate the head exploratory behavior of *Caenorhabditis elegans*, a minimal system capable of intricate motor patterns. Using variational mode decomposition, we identified two distinct motor dynamics: slow rhythmic bends propagating along the body and fast, phase-specific head casts influencing directional bias. Combinatorial ablations of three types of cholinergic motor neurons, in conjunction with dynamical systems analysis, revealed their distinct and overlapping roles: RMD contributes to head casts, SMD sustains bending states, SMB and SMD enable slow rhythmic bending and head-body coupling. Collectively, these neurons form a central pattern generator (CPG) driving forward locomotion. In addition, RMD subgroups displayed unique calcium dynamics correlated with rapid head movements and slow transitions between behavior states. We propose that dual-proprioceptive feedback underlies these multi-timescale dynamics, with slow feedback coordinating rhythmic bending and fast feedback shaping head casts, thereby optimizing roaming efficiency. These findings highlight how intrinsic dynamics and structured composition emerge from highly interactive low-level circuits.

## Introduction

Motor control in animals depends on creating highly structured and yet flexible movement patterns. Traditional theories emphasize hierarchical descending control, where higher-order brain regions command fixed sequences executed by lower-level circuits (1–4). This framework has been instrumental in explaining stereotyped movements such as rhythmic gaits (5–8). However, it falls short in accounting for the variability and adaptability inherent in more naturalistic and exploratory behaviors: for example, when a monkey is performing a reaching task (9), elephants manipulate their trunks to investigate and interact with their environment (10), and humans use quick eye movements to visually scan their surroundings (11). Recent findings suggest that motor dynamics involve the intrinsic properties of local circuits (9, 12–14), which integrate proprioceptive and environmental feedback on various timescales (15, 16), allowing flexible and coordinated movements. How do local circuits generate intrinsic dynamics and how do these dynamics facilitate efficient navigation and exploration?

The one-millimeter long nematode *Caenorhabditis elegans* is a compelling model for exploring these questions. In its microscopic world, *C. elegans* uses biased random walks (17) and weathervane strategies (18–20) to locate food and resources by balancing exploration and exploitation (21–23). Little is known about the fine and rapid head swings during navigation (24). Most sensory neurons in *C. elegans* expose their cilia ends at the tip of the head, acting as tiny antennae. Each head swing converts spatial cues into temporal codes to guide decisions on food, mates, and threats. These swings involve complex dynamics: fast movements (“casts” (25)) may enable quick environmental sampling and directional changes, while slower bends aid extended movement and coordination. Such fast and slow patterns are reminiscent of quick and fine stepping adjustments needed during walking and running. However, the mechanism and function of this duality are elusive.

The ventral-dorsal head movement in *C. elegans* is mainly driven by three classes of cholinergic motor neurons: RMD, SMD, and SMB. RMD includes six neurons, while SMD and SMB have four each (Fig. 1B). Ventral head muscles are innervated by RMDV, RMDR, SMDV, and SMBV; dorsal muscles by RMDD, RMDL, SMDD, and SMBD (24, 26–28). RMDV and SMDV have inhibitory connections with their contralateral counterparts. SMBV and SMBD do not directly connect but are linked indirectly by SAA interneurons (24, 29). The neurons are interconnected either by SAA or gap junctions (Fig. 1B). These connectivity patterns, featuring reciprocal inhibition, find parallels in B-type or A-type motor neurons in the *C. elegans* ventral nerve cord (30–32) and more broadly in many other motor systems as part of a central pattern generator (CPG) (6, 8, 33–36). Rhythmic head motion helps establish the body undulation patterns necessary for forward movement (6, 32, 34) and guides animals towards food sources through ventral-dorsal bending bias (37–41).

**Figure 1.**
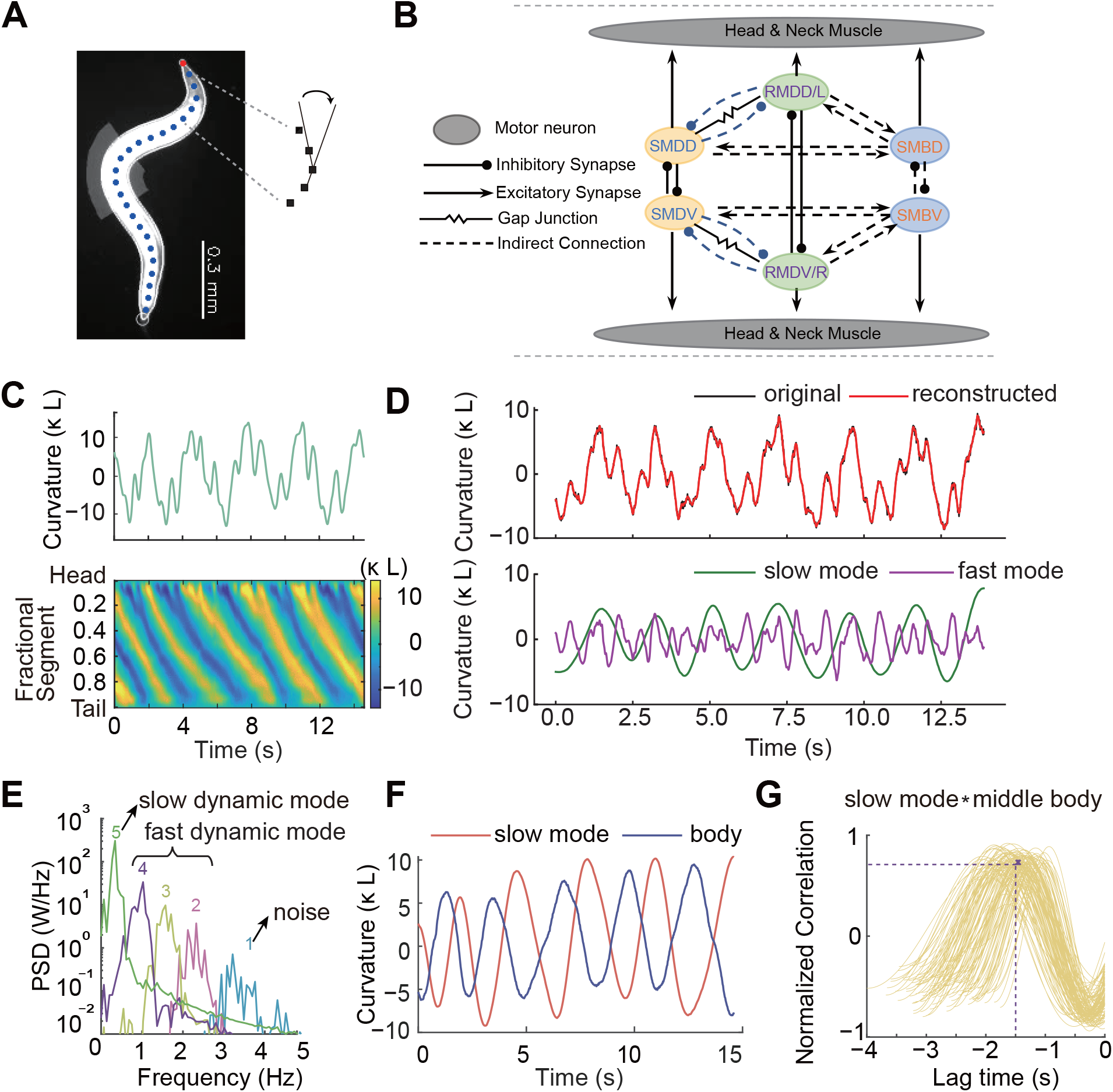
Slow and fast components of head movements. **A.** Centerline extraction (blue dotted line) for measuring body curvature along the worm’s length. **B**. Schematic of the putative head CPG circuit, consisting of SMD, RMD, and SMB motor neurons. Dashed lines indicate indirect connections bridged by SAA interneurons (not shown). **C**. Representative head curvature and body curvature kymograph of wild-type N2. *κ* and L denote the curvature variable and the worm length, respectively. **D**. Variational Mode Decomposition (VMD) of a representative head curvature signal. The reconstructed signal is obtained by subtracting the noise mode (mode 1) from the original signal. Slow dynamic mode and fast dynamic mode (slow mode and fast mode for short) correspond to the 5th mode and the summation of modes 2-4, respectively. **E**. Power spectral density (PSD) of each decomposed mode. **F**. Example time series of slow mode and middle body curvature (mean curvature of 45% - 60% portion of the body). **G**. Cross-correlation shows high peak correlation between slow dynamic mode and middle body curvature.

These findings raise interesting questions. If the goal of the head motor circuit is to generate rhythmic activity, why use *three* different types of motor neurons instead of one? Do they have unique or overlapping functions? The circuit motif elucidated by the worm connectome (28) and the head-bending correlated Ca^2+^ activity noted in certain neurons (25, 39, 42–44) do not readily provide clarification. By integrating experimental observations and computational analysis, we demonstrated that each of the three classes of cholinergic motor neurons exerts control over distinct aspects of head movements, working in synergy to create complex and structured dynamics that span multiple timescales. We propose a dual proprioceptive feedback framework to reproduce these movement patterns and demonstrate their role in optimizing movement speed. Therefore, instead of generating fixed motor sequences from top-down commands, efficient and adaptive movements emerge from highly interconnected and interactive low-level circuits that adeptly respond to real-time sensory feedback.

## Results

### Temporal decomposition of head movements

We recorded *C. elegans* spontaneous crawling behavior at a frame rate of ∼ 90 Hz using dark-field microscopy and computed the head curvature *κ*(*t*) during forward locomotion by averaging over the first 15% of the worm body centerline (Fig. 1A and Methods). To characterize the dynamics of complex head movements (Fig. 1C top), we applied variational mode decomposition (VMD) (45) to decompose the time-varying signal into different modes with decreasing central frequencies (Fig. 1D, E, Fig. S1A). A slower mode exhibits a higher magnitude in the power spectrum (Fig. 1E). The first mode, which entails the highest frequency (Fig. S1B), is noise. The sum of the remaining four modes (mode 2-5) is sufficient for an accurate reconstruction of head bending dynamics (Fig. 1D).

As illustrated in the spatiotemporal curvature kymograph (Fig. 1C bottom), only the slow rhythmic bending activity of the head back-propagated along the body during *C. elegans* forward locomotion (Fig. 1F). We therefore used VMD to identify the *slow dynamic mode κ*_*s*_(*t*) (e.g., mode 5) that has the highest cross-correlation with body bending activity (Fig. 1G and Methods). The sum of the remaining modes (e.g. mode 2-4) was considered a *fast dynamic mode κ*_*f*_ (*t*) intrinsic to the motor activity of the head (Fig. 1D bottom).

### Phase-dependent head casts promote shallow turns

Head curvature is sinusoidal, with distinct dips and peaks contributing to the fast dynamic mode. These rapid and intricate movements, reminiscent of the previously studied “head casts” (25), involve head bends that propagate and terminate before the middle body. We pinpoint the location of head casts, characterized by small and rapid bidirectional changes in head curvatures (Fig. 2A shaded). These unique events were identified in the time derivative of head curvature, since 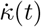 rapidly crosses zero twice, leading to a pronounced peak in the time series (Fig. 2A bottom and Methods). The larger amplitude head casts tended to propagate further along the body (Fig. S2A, B), while most had a limited propagating distance (Fig. S2C). We found that the occurrence of head casts exhibited a phase-dependent distribution (Fig. 2C and Methods) when aligned with the slow dynamic mode. Without loss of generality, we define *ϕ* = 0 when the slow-mode curvature *κ*_*s*_ is at its maximum and *ϕ* = *π* when *κ*_*s*_ is at its minimum. Consistent with Kaplan’s work (25), head casts occurred with a higher probability when *ϕ* was in the first or third quadrant (Fig. 2C).

**Figure 2.**
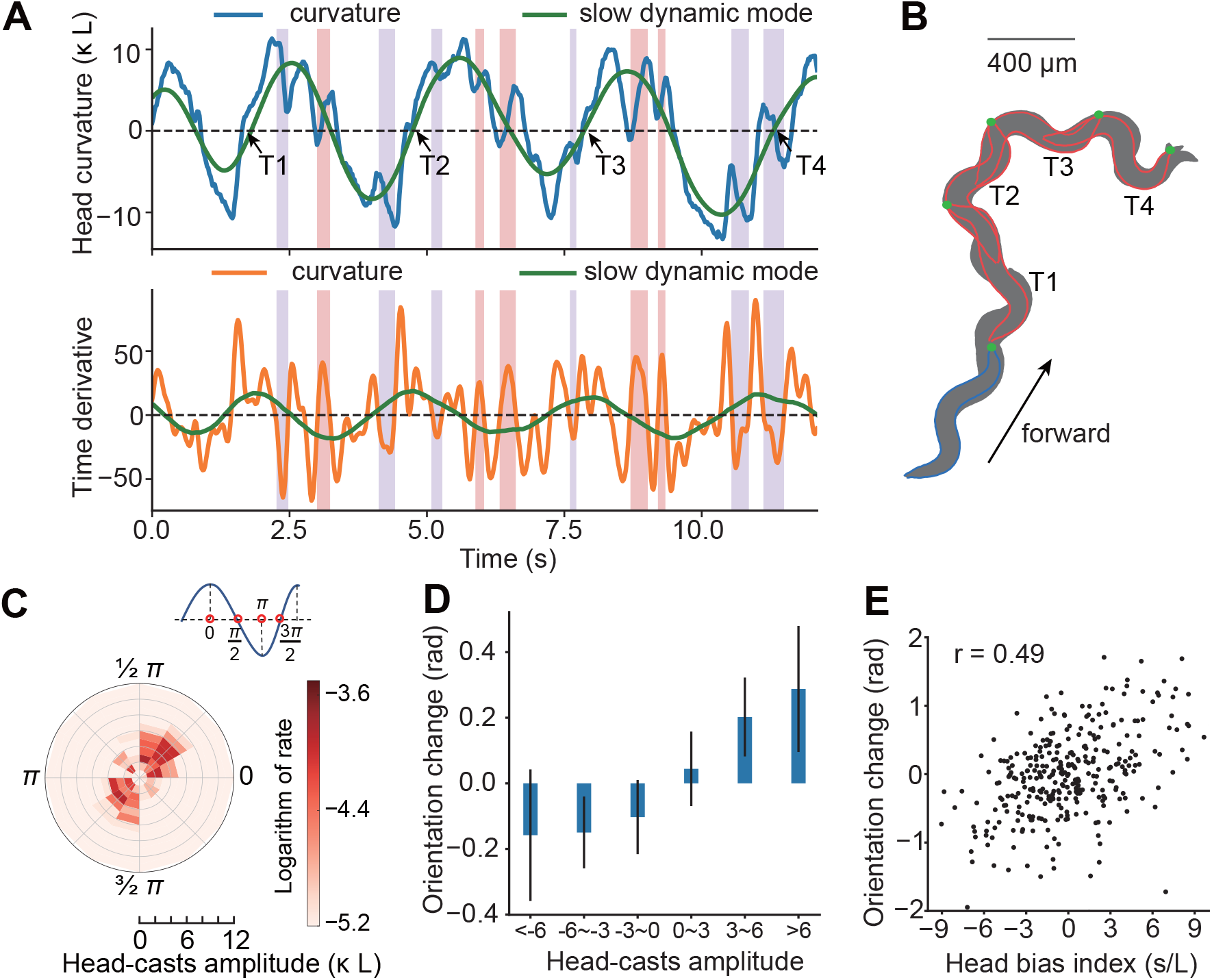
Illustration of head casts and their association with gradual directional changes. **A.** Top: Time series of the head bending curvature and the corresponding slow dynamic mode. The light red and light blue shaded regions correspond to the dorsal and ventral head casts, respectively. Bottom: Time derivatives of the denoised head curvature and slow dynamic mode. **B**. Sequential snapshots of the nematode’s posture over three consecutive undulation periods demonstrate a potential link between gradual directional changes and persistent dorsal head casting. **C**. The joint distribution of head casts across stroking amplitude and phase, where the phase is defined by the slow dynamic mode: 0 represents maximum dorsal bending and *π* indicates maximum ventral bending. The color map depicts the logarithmic head cast rate. **D**. A linear model is employed to assess the impact of head casts at different amplitudes on the directional changes during forward locomotion (Methods). Head casts were grouped into six amplitude categories: <-6, -6 to -3, -3 to 0, 0 to 3, 3 to 6, and >6, with negative levels representing head casts during negative curvature and positive levels during positive curvature. For each three-cycle period, we recorded the orientation change and number of head casts from six amplitude categories. Coefficients were determined using least squares regression. To quantify coefficient uncertainty, the datasets were bootstrapped 10,000 times, with standard deviations depicted as black error bars. **E**. The total orientation change and dorsoventral bending bias of the head (area under the curvature curve, Methods) were recorded over intervals of three head bending cycles (black dots). Pearson correlation coefficient *r* = 0.49, p = 3.5 *×* 10^−7^.

We sought to understand how phase-dependent head casts could influence exploratory behaviors on an extended timescale. We analyzed all head casts within three undulation cycles, using a linear regression model to predict changes in worm orientation during that period of movement (Methods). Linear regression revealed that a large amplitude head cast made a significant contribution to a change in movement direction (Fig. 2D). Importantly, these phase-dependent head casts extended the head bending duration towards one side (Fig. 2A), resulting in a ventral/dorsal bias of head bending. This led us to hypothesize that this bias might translate into changes in running direction (see example trajectory in Fig. 2B). To empirically assess this idea, we introduced a head bias index, calculated as the area under the curve *κ*(*t*) over three undulation cycles. Our subsequent analysis confirmed that this bias index is significantly positively correlated with the change in movement orientation (Fig. 2E) as this bias can propagate along the body (Fig. S2 F). Together, our findings underscore the importance of fine head movements in guiding exploratory behavior.

### Different cholinergic motor neuron classes contribute to distinct aspects of head bending dynamics

We sought to discern the roles of each type of cholinergic motor neuron in head movement using combinatorial optogenetics. With cell-specific promoters, we generated transgenic animals that allow optogenetic ablation of RMD, SMD, and SMB motor neurons either individually or in combination. We then quantified the changes in head bending dynamics *κ*(*t*) (Fig. 3A, Fig. S3A) by calculating the power of the slow dynamic mode *P*_*s*_ and the fast dynamic mode *P*_*f*_ (Fig. 1D and Methods). Interestingly, RMD-ablated animals saw a decrease in the contribution of the fast dynamic mode to head movements, calculated by the power ratio *P*_*f*_ */*(*P*_*s*_ + *P*_*f*_), whereas the contribution of the slow mode grew (Fig. 3B). However, SMD or SMB ablation reversed this trend, diminishing the slow-mode contribution.

**Figure 3.**
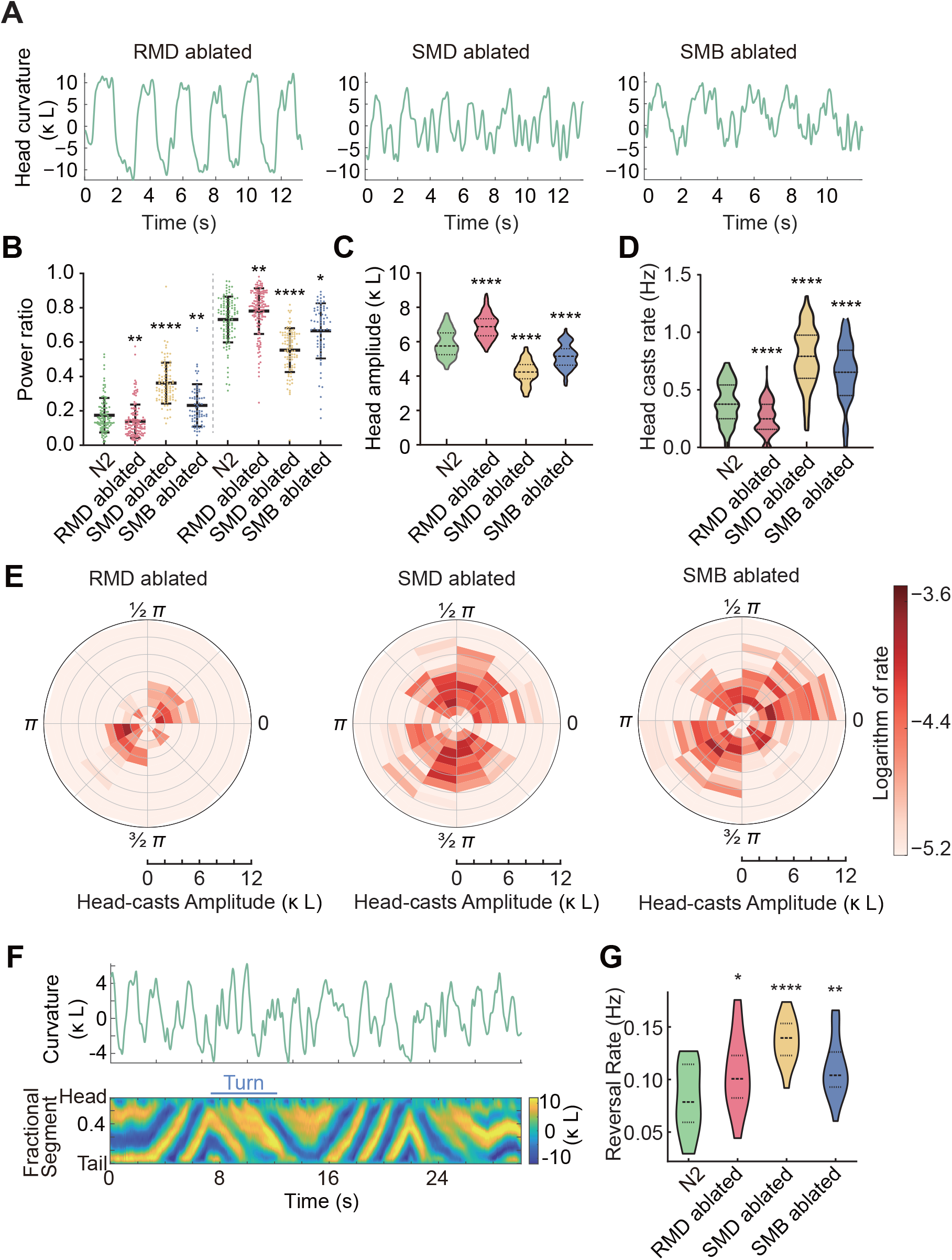
Ablation of different motor neuron types. **A.** Representative head curvatures in head motor neuron ablated animals. **B**. Power ratio of fast dynamic mode and slow dynamic mode in N2 (n=108, 18 worms), RMD-ablated animals (n=138, 19 worms), SMD-ablated animals (n=101, 15 worms), and SMB-ablated animals (n=85, 21 worms). n represents the number of episodes of forward movement. **C**. Head bending amplitude in N2 and motor neuron ablated animals. **D**. Head cast rate in each episode of forward movements. The rate is defined as the number of head casts with a large relative amplitude (Methods) divided by the duration of a forward run. **E**. The joint distributions of head casts over their stroking amplitude and phase in RMD, SMD, and SMB ablated animals. **F**. Example head curvature time series (top), and whole body curvature kymograph (bottom) from an RMD-SMD-SMB triple-ablated worm. **G**. Increased reversal rate for RMD (n=19), SMD (n=15), and SMB (n=21) ablated worms in their roaming state. For N2, n=18. n represents the number of worms tracked in long term. *p < 0.05, **p < 0.01, ***p < 0.001, ****p < 0.0001, Mann-Whitney U test. All tests are between wild-type N2 and mutants.

In examining the factors causing the power shift, we found that RMD-ablated animals displayed a higher head bending amplitude (Fig. 3C, Methods) but a lower head-cast rate (Fig. 3D) and a lower average head-cast amplitude (Fig. 3E left). On the contrary, SMD or SMB-ablated animals had a decreased head bending amplitude (Fig. 3C) and an elevated head cast rate (Fig. 3D, Fig. S3B). The changes in these parameters aligned with the observed shifts in power. Consistently, optogenetic inhibition of SMD suppressed the overall head/neck bending (Fig. S6A,B) and increased the contribution of the fast mode to the head bending power (Fig. S6C). Notably, while RMD ablation left the head cast’s phase dependence unaltered (Fig. 3E left), SMD or SMB ablation disrupted it, resulting in head casts distributed randomly across all phases of the slow dynamic mode (Fig. 3E middle and right).

Furthermore, in SMD- or SMB-ablated animals, the head-body coupling was disrupted. This is evident as the maximum cross-correlation between the slow dynamic mode and the body-bending activity was significantly reduced (Fig. S3C). Intriguingly, when RMD, SMD and SMB were co-ablated, the animals could not move forward, but backward movements remained unaffected (Fig. 3F). Our results suggest that continuous *C. elegans* forward locomotion requires sustained rhythmic bending activity initiated from the head motor circuit.

### Phase space analysis of head bending dynamics

Next, we applied time-delay embedding to reconstruct the multidimensional phase space from the one-dimensional time series *κ*(*t*) (Methods). According to Takens’ theorem, the structure of the reconstructed phase space would be topologically equivalent to the original system (46). This invariance offers a geometric perspective to deepen our understanding of the intricacies of head motor circuit dynamics.

We found that the phase trajectories of head movements could be largely embedded in 3 dimensions. The phase space coordinates are a linear transformation of the time-delayed *κ*(*t*), and thus are correlated with the head curvature and its time derivative (Methods). We used a contour density map and its maximum intensity projection to indicate how often local regions in the phase space were visited (Fig. 4). For wild-type animals, these phase-space trajectories appear circular, indicating periodic motion (Fig. 4A1). However, the irregularity in head movements causes localized scattering, making the overall phase space resemble a doughnut when visualized using a density plot.

**Figure 4.**
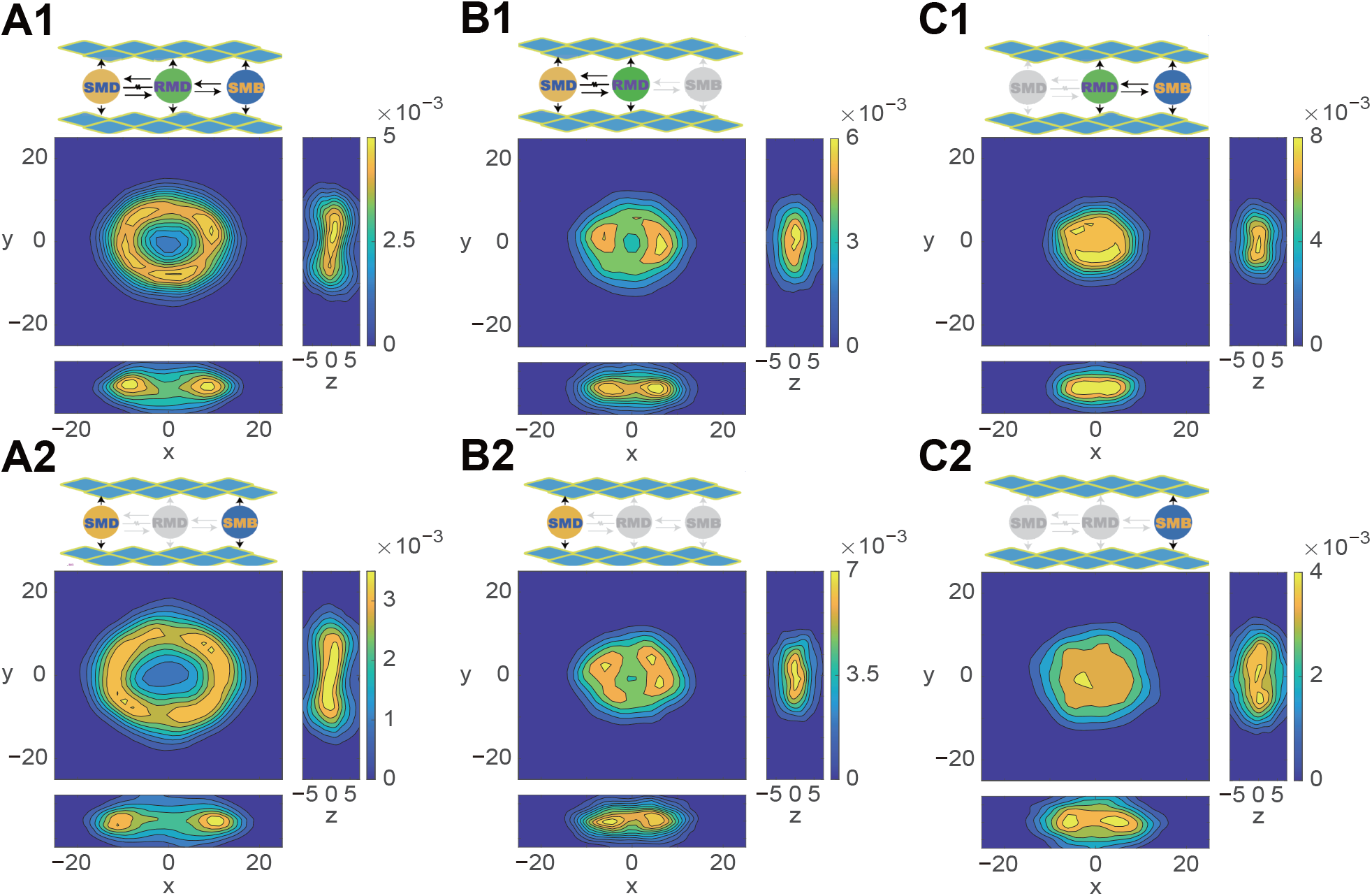
Exploratory head dynamics in an embedding phase space. Each animal’s head curvature time series during forward locomotion was embedded into a high-dimensional space with automatically estimated dimensions and delays. The trajectories were then projected onto three principal components (x, y, and z axes), forming a 3D embedding space. Color maps represent trajectory density estimation. Different motor neuron dominance led to varied phase space density features. At the top of each panel is a schematic of the ablation scheme with gray ellipses representing ablated motor neuron types. The second row shows figures with RMD ablated, and the second and third columns with SMB and SMD ablated. Topological features vary significantly across columns but show similarities between rows, reflecting dynamic characteristics: **A1-A2**. SMD-SMB dominated control shows a doughnut-shaped density, suggestive of limit cycle dynamics. **B1-B2**. Worms with SMB-ablated and SMD-dominated bending exhibit two symmetric density clusters, indicating bistable dynamics. **C1-C2**. SMB-dominant control displays a uniform disk-shaped density, indicating chaotic head bending.

We explored the influence of different motor neuron types on the topological features of head dynamics by conducting single and combinatorial ablation experiments on three types of motor neurons. This approach allowed us to observe changes in topology when head movements were governed by one or two types of motor neurons (Fig. 4). When governed predominantly by SMD (with and without RMD in Fig. 4B1 and B2, respectively), the phase space density displayed two nearly symmetrical clusters, suggesting bistable dynamics with the head in either a dorsal or ventral bending state. In head movements predominantly governed by SMB (with and without RMD in Fig. 4C1 and C2, respectively), the ring structure (in Fig. 4A1) collapsed, and the phase trajectories spread almost uniformly within a disc. Here, the phase space’s center, corresponding to minimal head bending velocity and amplitude, was visited more frequently than in wild-type animals. Interestingly, RMD ablation expanded the circular density distribution *without* changing its topology (Fig. 4A2), whereas SMD or SMB ablation significantly altered the topology. Together, SMD and SMB enable rhythmic head bending, while RMD neurons aid in fast head casts during the main oscillation.

### RMD subgroups display distinct and multi-timescale dynamics

Next, we conducted calcium imaging of RMD neurons (Fig. 5A) in freely moving animals. We further improved spatial resolution with high NA imaging in a microfluidic device that immobilized the worm’s body while allowing head movement (Fig. 5B). The activity of RMDL and RMDR remained indistinguishable due to their spatial overlap, and we thus use RMDL/R to represent either neuron. Our experiments reveal distinct activity patterns in RMD neuron subgroups across different behavior timescales. First, RMDD and RMDV neurons showed coordinated activity changes, with calcium levels ramping up simultaneously during forward-to-reversal transitions and decreasing synchronously during reversal-to-forward transitions (Fig. 5A bottom and Fig. 5C). During reversals, the head exhibited regular undulation with a significantly reduced frequency of head casts (Fig. 5E). Second, we found a strong correlation between the time derivative of the RMDL/R activity and that of the head bending signal (Fig. 5D). In contrast, RMDD/V neurons displayed a much weaker correlation (Fig. 5E). Taken together, these findings suggest that RMDL/R neurons directly drive rapid head bending, whereas RMDD/V neurons influence overall head movement patterns during forward and reversal states.

**Figure 5.**
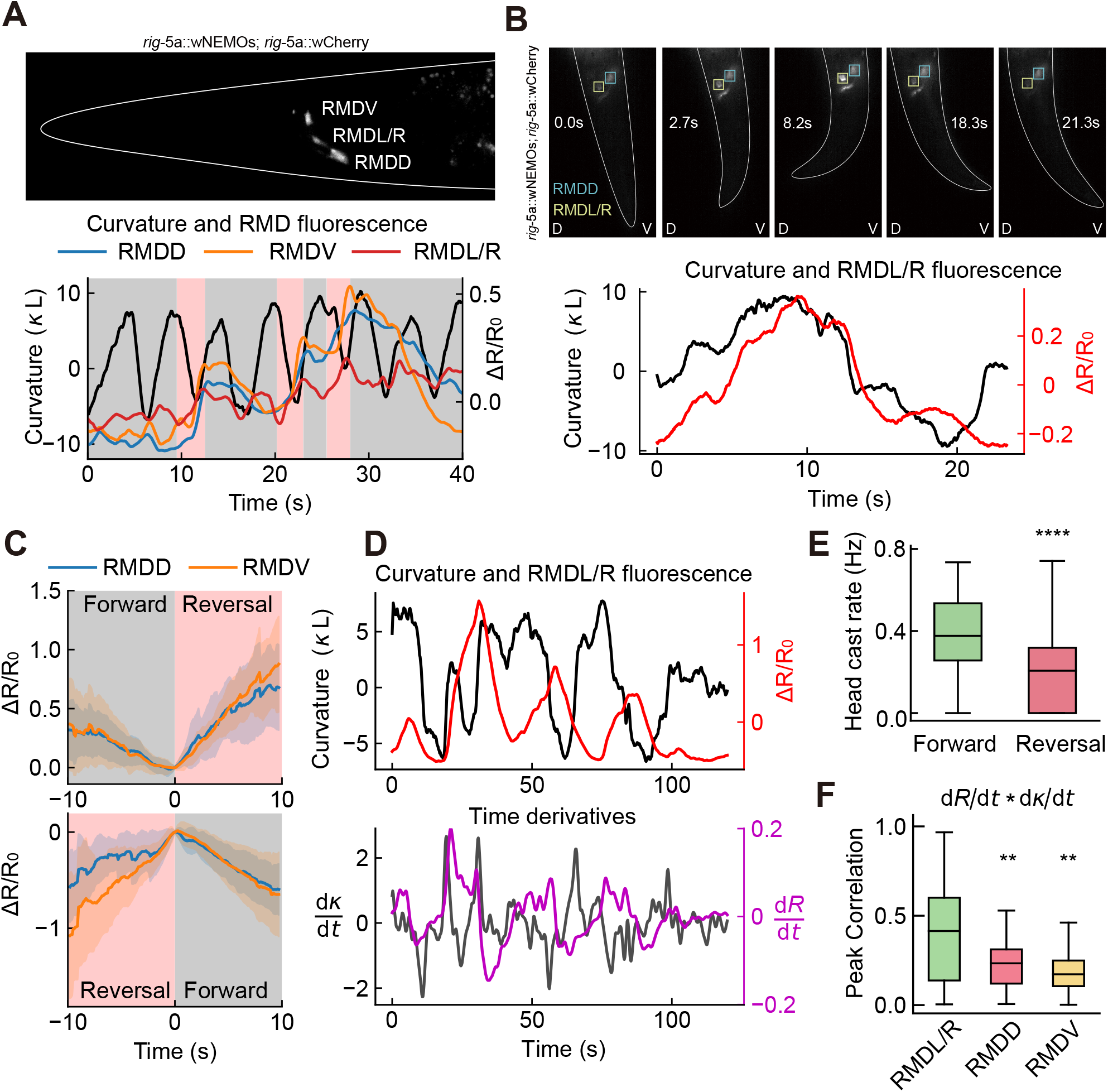
Calcium imaging of RMD neurons. **A.** Top: Fluorescent imaging of RMD neurons’ calcium activity. A representative image from high NA imaging in a microfluidic device. Bottom: Illustrative RMDD, RMDV, and RMDL/R activity in free movement imaging setting, comprising forward movement (gray background) and reversal episodes (red background). **B**. Top: Time-lapse fluorescent imaging of RMD neurons in worms with free heads but restrained bodies in microfluidic tunnels. Images show the activity of RMDL/R neurons during head movement at different time points (0s to 21.3s). D and V labels indicate dorsal and ventral sides; Bottom: RMDL/R activity and curvature increased and decreased synchronously. **C**. RMDD and RMDV activity ramped up together during forward-to-reversal transitions and decreased synchronously during reversal-to-forward transitions. Solid lines represent means of aligned fluorescent signals. Shaded areas depict standard deviations. **D**. Representative head curvature and RMDL/R activity and their time derivatives. **E**. Worms showed reduced head casting during reversals (n=77, 18 worms) than forward episodes (n=108, 18 worms). n represents the number of episodes of forward movement or reversal. **F**. The time derivative of head curvature (d*κ*/dt) correlates more closely with the time derivative of neural activities (dR/dt) of RMDL/R (n=42, 2 worms) than with RMDD (n=26, 3 worms) and RMDV (n=26, 3 worms). n represents the number of 10-second segments of time series. **p < 0.01, ****p < 0.0001, Mann-Whitney U test.

### Head casts enhance kinematic efficiency

For long-distance exploration, *C. elegans* must maintain an efficient crawling posture. Previous studies (47, 48) suggested that the angle of attack of crawling worms is close to an optimal value for maximum locomotion speed. Here, we provide a theoretical derivation considering the wavelength correction under large amplitude. Our derivation is based on linear resistive force theory, which effectively and accurately links the body posture kinematics to locomotion (49–52). The theory shows that the locomotion velocity is given by (Methods):

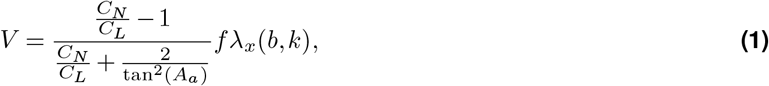

where *C*_*N*_ is the frictional drag coefficient normal to the body centerline, *C*_*L*_ is the one parallel to it, and *f* is the undulation frequency. *λ*_*x*_ is the effective wavelength along the movement direction: it decreases as the wave number *k*, the wave amplitude *b*, and the angle of attack *A*_*a*_ increase. Eq. (1) shows a small angle of attack increases the slip rate, the ratio between the wave velocity *fλ*_*x*_ and the locomotion velocity *V*, while a large angle reduces the effective wavelength. Both extremes negatively impact speed. Consequently, for a given wave number, there is an optimal angle of attack that maximizes the *kinematic efficiency*, defined as *V/f*. This efficiency measures the distance traveled per undulation cycle and is independent of the undulation frequency (Fig. 6B). Our analysis showed that RMD-ablated animals displayed an increase in head and body curvature (Fig. 6A bottom, C) and the angle of attack (Fig. 6D top), along with a decrease in locomotion velocity and kinematic efficiency (Fig. 6D middle and bottom).

**Figure 6.**
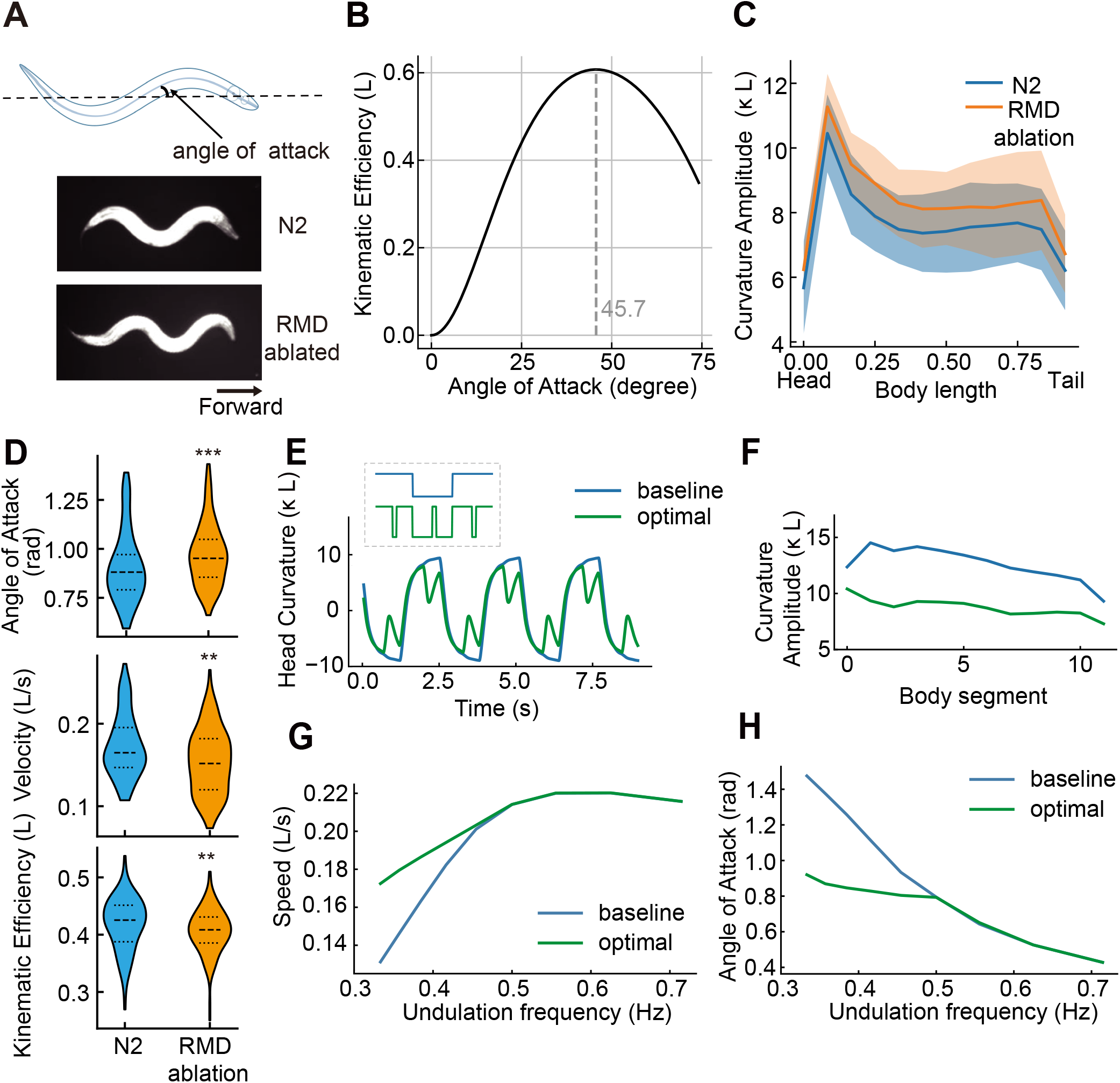
Locomotion Efficiency. **A.** Angle of attack represents the average angle between the body and movement direction (top). Wild-type N2 (middle) and RMD-ablated worms (bottom) exhibit distinct postures during forward locomotion. **B**. A non-monotonic relationship between kinematic efficiency and angle of attack indicates an optimal angle of attack around 45 for maximum locomotion efficiency. The ratio of drag coefficients *C*_*N*_ */C*_*L*_ = 10 follows recent estimations on agar surface (53–55). **C**. Optogenetic ablation of RMD neurons increased overall body curvature. **D**. RMD-ablated worms (n=138, 19 worms) demonstrate increased angle of attack (top, p = 7.5 *×* 10^−4^), decreased crawling speed (middle, p = 7.1 *×* 10^−4^), and reduced kinematic efficiency (bottom, p = 8.9 *×* 10^−3^) compared to N2 controls (n=108, 18 worms); n denotes the number of episodes of forward run. **p < 0.01, ***p < 0.001, Mann-Whitney U test. **E**. Comparison of head curvatures driven by a baseline square wave input and the optimized head muscle input (inset, dashed box). The head input period is fixed at 2.4 s. **F**. Body curvature along the worm’s length shows an overall decrease following the optimized head input from **E. G-H**. Locomotion speed and angle of attack under baseline and optimized head input across various undulation frequencies.

Given the worm’s ability to bend into large curvatures, how does the neural circuit control the amplitude for efficient crawling? What is the optimal head muscle control signal? To this end, we assume that the head neural circuit produces a binary input signal, alternating between dorsal and ventral head muscle activation. We simulated head bending in response to control signals based on a realistic mechanical model and propagated this bending along the body (Methods, (32, 50)). We integrated fast switches in a slow periodic input signal and optimized the pattern to maximize crawling speed. The optimal head curvature resembles the observed phase-specific head casts nested in a slow oscillation (Fig. 6E). These simulated head casts reduced both head bending amplitude and overall body curvature (Fig. 6F) as the bends propagated backward. This optimized movement pattern increased locomotion velocity by 14%, consistent with our theoretical predictions. Reversing the phase of the fast switches (Fig. S5D) led to larger head bending, similar body bending curvatures (Fig. S5E) and a slight decrease in velocity. Thus, our optimization method clarifies how head casts improve locomotion efficiency and partially explains their timing. The head motor circuit is likely to impose additional constraints that affect the phase dependency of head casts.

The head cast strategy boosts speed by lowering the average angle of attack at frequencies below 0.5 Hz and aligns with the baseline square-wave input above 0.5 Hz (Fig. 6G, H). Our model thus predicts that when the head oscillation becomes sufficiently fast, head cast will vanish, which agrees with a previous finding (25). Furthermore, our model predicts that the use of the head-cast reduces muscle energy expenditure due to a decrease in overall curvature (Fig. S5F). Together, our results suggest that by optimizing head muscle input in our neuromechanical model, phase-specific head casts effectively control crawling posture, maximizing locomotion velocity and minimizing energy use during exploration.

### A bend-sensitive feedback model captures slow and fast dynamics

We asked whether the observed fast and slow head-bending activity could be reproduced by a minimal model that is consistent with known anatomical and functional data. Our ablation experiments demonstrated the synergistic role of SMD and SMB as a core oscillator and the role of RMD in the generation of head casts. Direct or indirect cross-inhibition between contralateral SMD/SMB neurons, as well as our ablation studies, suggest bistable switches in the slow oscillator. We modeled SMDD and SMBD as one dorsal unit and SMDV and SMBV as one ventral unit (Fig. 7A left), activating dorsal and ventral muscles, respectively. Both units exhibit bistable dynamics, modeled as the hysteresis curve (Fig. 7A right, (56)). SMD and SMB, with posterior sublateral processes, are hypothesized to sense posterior curvature. Thus, our model includes proprioceptive feedback from neck bending to the core oscillator, acting as a slow integration of head curvature, captured by *K*^slow^ (Fig. 7A left, Methods).

**Figure 7.**
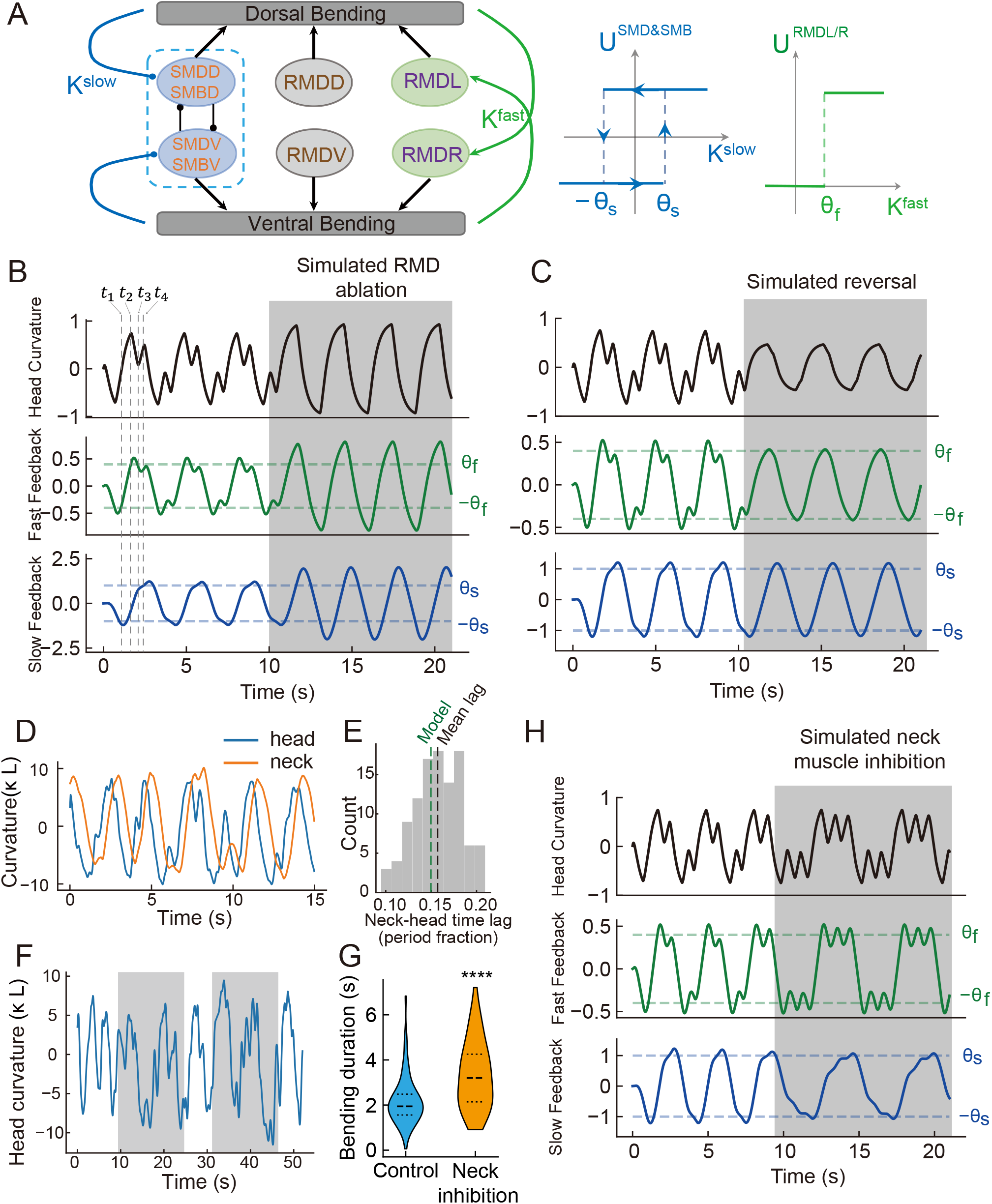
A circuit model captures the fast and slow dynamic modes. **A**. Schematic of the model circuit and the bistable dynamics of its elements. SMD and SMB motor neurons on the same side operate as one functional unit and receive a slow integration of the head curvature *K*^slow^ as feedback. The mutually inhibiting dorsal-ventral units were posited to operate in a bistable regime and have a hysteresis curve in the input-voltage relation (middle). RMDL and RMDR activate the dorsal and ventral muscles, respectively. Their activation levels are switched by a fast integration of head curvature *K*^fast^ (right). RMDD and RMDV provide behavior state-dependent input: they remain silent during forward and provide identical inputs to muscles when simulating reversal. **B**. Representative trajectories of head curvature, fast feedback, and slow feedback. Dashed lines indicate the four different circuit states: *t*_1_: dorsal SMD-SMB unit was activated. *t*_2_: The fast curvature feedback activated RMDV/R, generating an opposite muscle torque. *t*_3_: decreased fast feedback shut RMDV/R down. *t*_4_: slow curvature feedback switched the core oscillator to ventral outputting. The shaded area simulated RMD inhibition, which showed increased head bending amplitude due to lack of fast feedback. **C**. Simulated head movement during forward movement and transiting to reversal (shaded period). **D**. Representative head (0-15% body length) and neck (15%-30% body length) curvature from wild-type animals. **E**. Histogram of time lags between head and neck curvature in N2 worms (n=108, 18 worms); The mean time lag was 0.16 undulation period, consistent with the time lag between head curvature and surrogate posterior curvature in the model (0.15 period). **F**. Representative head curvature during optogenetic neck inhibition. Illumination (shaded area) induces extended bending toward both sides. **G**. Significant increase in time spent bending toward both sides during neck inhibition (n=50, 2 worms). For control: n =75, 2 worms; p = 1.1 *×* 10^−5^, Mann-Whitney U test. **H**. Simulated neck inhibition (shaded area) showed excessive head casts and extended bending duration.

On the fast timescale, RMDL/R motor neurons activate dorsal and ventral muscles respectively, operating in a bistable state according to electrophysiological measurements (57). As upstream mechanosensitive neurons (e.g., URY, IL1, OLQ, and CEP) may sense head pressure (24, 58, 59), we hypothesize that RMDL/R receive rapid feedback from head curvature, captured by *K*^fast^, toggling neural activity on and off (Fig. 7A right). Our model thus includes two parallel neuronal groups and a dual feedback system that integrates head curvature over two distinct timescales. In addition, we propose that RMDD/V functions on the timescale of behavioral states to influence head cast frequency (Fig. 5E). In reversal, coordinated RMDD and RMDV activation (Fig. 5C) may reduce head casts by synchronizing dorsal and ventral head muscle activity. In the model, these neurons remain silent during forward movement and provide identical input to head muscles when simulating head movement in reversal.

When the head bends dorsally (positive curvature, Fig. 7B, *t*_1_), fast feedback follows the head curvature and activates the RMDL neurons at the threshold *θ*_1_ (at *t*_2_ in Fig. 7B), producing opposite muscle torque and reducing head curvature. The fast feedback signal then drops below *θ*_1_, turning off the RMDR neurons (at *t*_3_ in Fig. 7B). However, the input to the dorsal muscles from the core oscillator persists, increasing the dorsal bending. Therefore, fast negative feedback could generate phase-specific head casts. The activity of the core oscillator is regulated by a low-pass filter of the head curvature *K*^*slow*^ that mimics the neck movement. Indeed, the observed phase lag between head and neck curvature in real animals closely matches the lag between head curvature and surrogate posterior curvature in our model (Fig. 7D, E, Methods). Sustained bending enhances this negative feedback, changing the neuronal state from dorsal to ventral activation (at *t*_4_ in Fig. 7B) and producing the slow dynamic mode. Ablating RMD neurons in simulations by removing fast negative feedback showed a similar increase in bending amplitude as seen in experiments (Fig. 7B, Fig. 3A). Furthermore, when switching from forward to reverse movement, RMDD and RMDV provide identical inputs to dorsal and ventral head muscles. This saturated activation reduces the activation difference between muscle groups, producing diminished torque for head bending and failing to trigger RMDL/R-mediated head casts (Fig. 7C).

A key prediction of the model is that the feedback originating from neck bending plays a key role in defining the slow dynamic mode. To test this, we reduced the neck curvature by optogenetically inhibiting the neck muscles. Upon inhibition, the head bent to one side for a long time, significantly increasing the duration on both sides (Fig. 7F,G). This observation can be captured by our model: reducing the amplitude of the slow feedback signal led to a longer accumulation and thus extended the period of the slow oscillator (Fig. 7H), consistent with the persistent head bending state observed in the experiments.

## Discussion

Bilaterians have thrived evolutionarily because their bilaterally symmetrical body plan enables advanced directed movement. This movement involves efficient forward propulsion and controlled turning, with head movement being a key to both maintaining forward speed and steering in animals like fish (60) and snakes (61). The *C. elegans* head motor circuit, a minimal system enabling highly adaptable movements, demonstrates motor patterns with a structured composition where rapid, phase-specific head casts are nested within slower head oscillations (25). Here we show that such intrinsic dynamics emerge from the interactions among three types of cholinergic motor neurons, each contributing a distinct role. SMD and SMB neurons together engender robust, limit-cycle-like slow head oscillations, whereas RMD exerts influence on head casts without modifying the overall topology of the dynamics (Fig. 4). During reversals, the dynamics of head bending exhibit increased regularity accompanied by a marked reduction in head casts (Fig. 5E and Fig. S7). Surprisingly, calcium imaging suggests unique functions within RMD neuronal subgroups: RMDL/R likely drives rapid head movements, whereas the transitions from forward to reversal motor states are characterized by elevated and synchronized calcium activity within the RMDD/V.

These findings collectively suggest that the coordinated activity of SMD, SMB, and RMD neurons underpins the hierarchical organization of head movements. Interestingly, in gap junction deficient mutants (*unc-7* ;*unc-9*), head casts are emitted randomly across different phases (Fig. S8), underscoring the importance of intercellular coupling in maintaining phase-dependent coordination. The regularization of dynamics during the reversal state implies a *shift* toward more stereotyped, feedforward descending control, with RMDD/V synchronization acting as a landmark for motor state transitions.

This modular architecture – where slow oscillations provide a scaffold for context-dependent, faster maneuvers – exemplifies a general principle for balancing robustness and flexibility in motor systems. Central pattern generators (CPGs) in the vertebrate spinal cord orchestrate slow, rhythmic movements like walking and swimming (36), analogous to the role of SMD/SMB neurons. Superimposed on these slow rhythms are faster, corrective adjustments mediated by sensory feedback (36), particularly proprioception, which bears a strong resemblance to the function of RMD neurons. Studies in *Drosophila* also reveal a hierarchical organization of leg motor neurons with distinct fast and slow types controlling different aspects of movement and exhibiting a recruitment order based on force production (62). This functional specialization in another invertebrate model provides additional evidence for the conservation of this strategy.

What is the behavioral significance of head casts? These rapid head movements propagate posteriorly along the body for a limited distance, and this observation remains consistent despite the differences in the parameters used to define the spatial extent of head curvature between the previous work (25) and ours (Fig. S2E and C). Here we discovered a significant correlation between head casts and worm reorientation, along with a causal relationship between dorsal-ventral bending biases in the head and the middle body where head bends propagate, linking intricate head movements with the worm’s overall turning behavior. Prior research has shown that the curved paths during weathervane behavior stem from asymmetric dorso-ventral modulation of head bending (63). Theoretical and empirical studies suggest that this occurs through differential gain modulation of contralateral head motor neurons in response to a chemical gradient (38, 64, 65). We propose head casting as a new navigation strategy, achieved by a dorso-ventrally asymmetric number of head casts. In essence, adjusting the timing and amplitude of head casts alters the phase and amplitude of the slow core oscillator. Future research should confirm the role of head casting in reorientation amid sensory signal gradients.

The head is crucial for leading forward movement, with its undulation necessary for coordinating body waves and propulsion. Head and body movements are closely coupled, largely through directional proprioceptive coupling (32), resulting in a strong correlation of amplitude and entrainment of the undulation frequency (6, 34). Here we demonstrated that RMD, SMD and SMB constitute the major component of the head CPG. When all three motor neuron types were ablated, *C. elegans* could not propel itself to move forward. Furthermore, using a computational model, we discovered that control signals for head casts emerged from optimizing movement velocity by tuning the overall bending amplitude close to an optimal value. This explains the ubiquity of head casts in the locomotion of *C. elegans*: they rely on this strategy to dynamically adjust their posture, keeping it optimized for efficient long-distance exploration.

In conclusion, we proposed a model for *C. elegans* exploratory head movements, where slow and fast dynamic modes emerge from the interplay between dual-timescale negative proprioceptive feedback and the bistable dynamics of the circuit components. Fast feedback to the RMD neurons likely comes from diverse mechanosensitive neurons, such as URY, IL1, OLQ, and CEP (24, 58, 59). Individual RMD neurons have a bistable membrane potential (57), a mode that allows nonlinear responses to mechanosensitive inputs. The neck and body act as low-pass filters, delaying and smoothing head movement. The posterior sublateral extensions of SMD and SMB neurons could act as stretch receptors, detecting the posterior curvature and providing slow feedback. This circuit modulates the bistable state transitions of the core oscillator to generate a slow dynamic mode. Together, rapid local feedback, delayed slow feedback, and neural input-output nonlinearity shape the dual oscillatory head movements and work collectively to optimize locomotion efficiency.

## ACKNOWLEDGEMENTS

The authors appreciate helpful discussions with Dmitri Chklovskii, Shangbang Gao, Yuichi Iino, Manuel Zimmer, and Mei Zhen. PL, HZ, YS, JW, LT and QW were partially supported by the Natural Science Foundation China through Major International (Regional) Joint Research Project (32020103007).

## Methods

### *C. elegans* strains

*C. elegans* strains were cultivated by standard procedures (66). Transgenic worms in optogenetic experiments were cultivated in the dark on NGM plates with OP50 bacteria and all-trans-retinal (ATR) for over 8 hr. We used young adult hermaphrodites to carry out all the experiments.

### Molecular biology

We used standard molecular biology methods. Promoters P*rig-5a* (2.2 kb), P*flp-22p-*Δ*4* (1.5 kb), and P*odr-2* were amplified by PCR from the N2 genome.

### Behaviour recording

We recorded *C. elegans* behaviours on 0.8% (wt/vol) M9 agar plate. Before recording, the worms were first transferred to a sterile NGM plate to eliminate the OP50 bacteria, then transferred to another sterile NGM plate for 25-35 min to make more frequent forward movements. The worms also acclimatized to the M9 agar for 2-3 min. Then we recorded freely moving worms for 5-15 min. Under dark field infrared illumination, the worms were automatically tracked and retained within the field of view of a 10 × objective on a Nikon inverted microscope (Ti-U, Japan). Behaviors were recorded by a Basler CMOS camera (aca2000-340kmNIR, Germany). We used MATLAB custom software (MathWorks, Inc Natick, MA, USA) to process the behavioral data afterwards.

### Optogenetic ablation

We used a mitochondrial-targeted miniSOG (TOMM20-miniSOG) (67) or a membrane-targeted miniSOG (PH-miniSOG) (68) to ablate specific neurons in *C. elegans*. Late L4/early young adult transgenic worms were transferred to an unseeded NGM plate, on which the worms were restricted by a 1 cm diameter filter paper ring soaked with 100 µM CuCl_2_. The worms were then illuminated for 5 min using pulse blue LED (M470L3-C5; Thorlabs) light (0.5 s on and 0.5 s off) at an intensity of 2.0 mW/mm^2^. After illumination, worms were transplanted to an OP50-seeded NGM plate and recovered over 4hrs before behaviour recording.

### Optogenetics

Lasers and a digital micromirror device (DLI4130 0.7 XGA, Digital Light Innovations, TX, USA) were used to generate spatiotemporal optogenetic manipulations (69) at a specific wavelength (473 nm, 561 nm or 635 nm). We manipulated the activities of neurons through light-activated channels (GtACR2, Arch, or Chrimson). Each worm was stimulated 5–8 times with at least a 40-second inter-stimulus interval.

### Behavioural analysis

We exclusively analyzed the forward movement. Forward-moving frames were manually labeled and further processed in MATLAB to quantify parameters. Forward movement was identified when the trajectory of a worm remained straight or turned no less than 90 degrees.

The worm orientation is the average angle between the x-axis of the camera frame and the 100 centerline segments. To quantify orientation changes during forward movement, the following steps were executed: 1. The curvature vectors of the centerlines were projected onto the first two principal components, producing two-dimensional postural representations denoted as *a*_1_(*t*) and *a*_2_(*t*). 2. The phase of forward movement (Φ) was computed with Φ = angle(*a*_1_ + *i* · *a*_2_), with zero-phase instances occurring when *a*_1_ is at its maximum and *a*_2_ is zero. 3. Orientation changes were measured between zero-phase moments separated by three cycles.

### Calcium imaging

Calcium imaging was performed in worms that expressed wNEMOs and wCherry proteins in the same neurons. The measurement of calcium activity was achieved by calculating a ratiometric change, which is defined as the fluorescence ratio of the green channel and the reference channel Δ*R*(*t*)*/R*_0_ (wNEMOs / wCherry), where *R*_0_ is the baseline ratio. Due to the broader emission spectrum of wNEMOs, which caused fluorescence bleed-through into the reference channel (wCherry), we corrected the activity ratio using the following equation

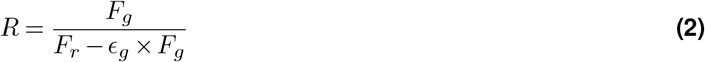

where *F*_*g*_ represents the fluorescence intensity of the green channel, *F*_*r*_ represents the fluorescence intensity of the reference channel, *ϵ*_*g*_ is the bleed-through coefficient of wNEMOs, which could be 0.08 to 0.20 under different experimental conditions.

To effectively capture calcium activity in RMD neurons during behavioral state transitions in freely moving worms crawling on a 2% (wt/vol) NGM agarose plate with a coverslip, calcium imaging was performed using a wide-field fluorescence microscope with a 20X objective (Nikon S Plan Fluor, WD = 7.4mm, NA = 0.45, Japan), which had a large imaging field (Fig. 5A bottom). Furthermore, to reduce motion artifacts and more accurately record calcium activity in the RMDL/R neuron during worm head swinging, we immobilized the worm body in a straight microfluidic channel where only the worm head can swing and used a 60X water immersion objective (Nikon Plan Apo, WD = 0.27 mm, NA = 1.2, Japan) instead of a 20X objective for higher resolution (Fig. 5A top, Fig. 5B and Fig. 5D). The objective was attached to a piezoelectric scanner (Physik Instrumente, Germany) to scan neurons along the z-axis. Blue light (488 nm) and green light (561 nm) were introduced to, respectively, excite wNEMOs and wCherry proteins. Green and red emission signals were captured by the objective, separated by a dichroic mirror, and projected onto sCMOS sensors (Andor Zyla 4.2, UK). The two-channel images were processed by custom-written MATLAB scripts.

### Statistical test

Statistical tests were indicated in the figure legends, including methods, error bars, numbers of trials and animals, and p-values. Using MATLAB, we applied the Mann-Whitney U test to determine the significance of the difference between the groups, and all multiple comparisons were adjusted using the Bonferroni correction.

### Kinematic analysis of head movements

We used custom-written MATLAB scripts to extract the timestamp, stage position and centerline of a worm. The centerline of the worm was first divided into *N* = 100 segments and the orientation of each segment, *θ*(*s*), *s* = 1, 2,···, *N*, was calculated. The curvature *K*(*s*) = Δ*θ*(*s*)*/*Δ*s* was calculated and then normalized into a dimensionless unit *K* · *L*, where *L* is the body length. And the dimensionless head curvature *κ* was calculated as

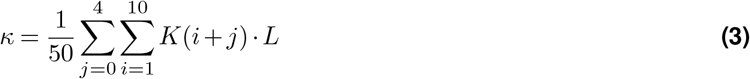

The head bending amplitude of each forward run episode was calculated using the following procedure: 1. A reference amplitude was introduced as *A*_ref_ = (*κ*(*t*)^Max^ − *κ*(*t*)^Min^)*/*2. 2. Using the scipy.signal package, we detected peaks in the curvature time series that exceeded the reference amplitude *A*_ref_ in prominence. These peaks correspond to maximum head-bending positions. The final amplitude was computed as the median of the absolute values of these identified peaks. The neck curvature is defined as the average curvature of the anterior 15%-30% region of the body and is calculated in the same way as the head curvature.

### Variational mode decomposition

We use the variational mode decomposition, abbreviated as VMD(45), to distinguish the fast and slow components of head bending dynamics. The head curvature signal was decomposed into five different intrinsic mode functions (*κ*_*i*_, *i* = 1, 2, 3, 4, 5) with decreasing central frequencies. The first mode *κ*_1_, which has the highest frequency, corresponds to the noise. The remaining four modes can reconstruct the fast and slow dynamics of head bending: a fast dynamic mode *κ*_*f*_ (*t*) is reconstructed by a linear combination of the modes 2-4, and a slow dynamic mode *κ*_*s*_(*t*) is identified by the mode 5.

The power of each mode *P*_*i*_ was calculated as 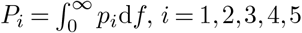, where *p*_*i*_ is the power spectrum density of mode i. Then, the power of the fast dynamic mode was calculated as *P*_*f*_ = *P*_2_ + *P*_3_ + *P*_4_, while the power of the slow dynamic mode was calculated as *P* = *P*. Subsequently, the power ratio was calculated as 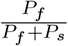 for the fast dynamic mode and 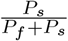 for the slow dynamic mode.

### Head-cast analysis

The rapid oscillation superimposed on the slow dynamic mode can sometimes propagate backward along the anterior body, a phenomenon known as the “head cast” (25). We extend this term to include smaller and rapid wiggles restricted to the very front part of the head.

Head-casts notably move opposite to the slow dynamic mode, identified in this study via a two-step process:

1. We identified zero-crossing points in the time derivative of a denoised head curvature time series 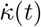, where *κ*(*t*) = *κ*^*s*^(*t*) + *κ*^*f*^ (*t*).
2. We distinguished these time points: zero crossings after which the time derivatives of the denoised signal and the slow dynamic mode have opposite signs 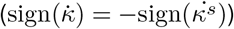 mark the start of a head cast (Fig. 2A). The next zero-crossing point is designated as the end of a head cast, after which 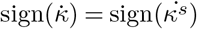.

The head-cast amplitude was measured as the absolute difference in head curvature between the start and end points. The relative amplitude, the ratio of absolute amplitude to head bending magnitude of the forward run, was used to identify significant head casts with relative amplitudes greater than 0.3, thus filtering out the head casts originating from the noise of curvature measurement. The occurrence rate was calculated as the number of significant head casts per episode divided by its duration.

We defined the phase of head bending using the slow dynamic mode, given its sinusoidal nature. Specifically, we used the instantaneous phase from its Hilbert transform.

### Linear regression to relate head cast and turning angle

We use a linear model to capture the relationship between the head casts and the gradual turning angle that the animal perform. Head casts were grouped into six amplitude categories: <-6, -6 to -3, -3 to 0, 0 to 3, 3 to 6, and >6, with negative levels representing head casts during negative curvature and positive levels during positive curvature. For each three-cycle period, we recorded the orientation change and number of head casts from six amplitude categories. The head casts were linearly aggregated to create an impact vector. if a single head cast with an amplitude <-6 and two head casts with amplitudes between 3-6 occur during a period, the resulting head-cast impact vector would be [1, 0, 0, 0, 2, 0]. To quantify the contribution of head cast from six amplitude categories, we run a linear regression to obtain the linear coefficients using least squares. To estimate the uncertainty in the coefficients, we performed bootstrap resampling, generating 10,000 randomized datasets and corresponding coefficient sets. This approach provides a robust estimation of the model’s parameter variability and statistical confidence.

### Phase space reconstruction

To analyze the dynamics of head movement, we employed phase space reconstruction using time-delay embedding, a technique based on Takens’ theorem (46). This method allows us to reconstruct the multidimensional phase space from our one-dimensional time series of head curvature *κ*(*t*).

To estimate the optimal time delay *τ* and embedding dimension *m*, we used MATLAB *phaseSpaceReconstruction* toolbox, which uses Average Mutual Information (AMI) (70) and False Nearest Neighbor (FNN) algorithm (71) to determine the delay time and embedding dimension respectively.

With *τ* and *m* determined, we constructed the phase space vector *y*(*t*) from the time series*κ*(*t*) as follows: *y*(*t*) = [*κ*(*t*), *κ*(*t* + *τ*), *κ*(*t* + 2*τ*), …, *κ*(*t* + (*m* − 1)*τ*)] This process transforms our one-dimensional time series into a set of m-dimensional vectors.

To visualize the topological feature of the reconstructed phase space, we projected the m-dimensional vectors into the space spanned by three principal components. For better visualization, we then fitted the density plots using kernel density estimation with a Gaussian kernel, whose optimal bandwidth was determined by the MATLAB *ksdensity* function.

This reconstructed phase space provides a geometric representation of the system’s dynamics, allowing us to identify features such as attractors, periodic orbits, or chaos, which are not apparent in the original time series.

### Circuit Model

The head circuit model consists of two parallel neuronal groups that innervate the head muscle: the RMD group and the SMD-SMB group. These groups utilize proprioceptive feedback from different time scales to modulate their states and output. The RMD group integrates rapid feedback from upstream mechanosensitive neurons, which are likely to detect head movements. Specifically, this fast feedback signal (*K*^fast^) is a low-pass filtered version of head curvature.

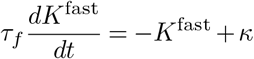

On the other hand, the SMD-SMB group receives slow feedback *K*^slow^ from posterior curvature, presumably detected by their posterior sublateral processes. This slow feedback signal *K*^slow^ is a low-pass filtered version of the surrogate posterior curvature −*κ*^*′*^, which is a slow integration of head curvature:

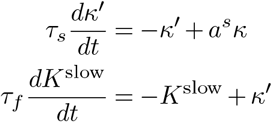

where the time constants are *τ*_*f*_ = 0.3*s* and *τ*_*s*_ = 0.8*s*.The parameter *a*^*s*^ determines the scale of surrogate posterior curvature *κ*^*′*^.

The bistable dynamics of the SMD-SMB group are represented using a hysteresis curve (Fig. 7A middle):

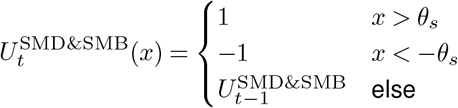

In this context, +1 represents dorsal muscle activation, while -1 represents ventral activation. For convenience, in the following equations, we denote the dorsal and ventral activation of the main oscillator as *U* ^SMDD&SMBD^, *U* ^SMDV&SMBV^, respectively:

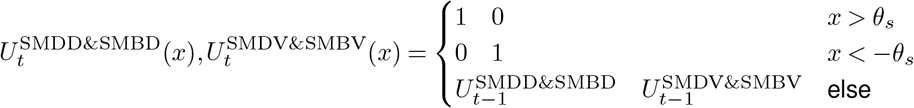

When the slow feedback exceeds *θ*_*s*_ or falls below −*θ*_*s*_, the neuronal output switches to the dorsal muscle activation (upper branch) or ventral activation (lower branch) in Fig. 7A middle, respectively.

The output voltage of the RMDL and RMDR neuron is modeled as a Heaviside step function of fast feedback.

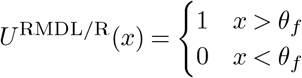

The output of RMDD and RMDV neurons, *U* ^RMDD^ and *U* ^RMDV^, are behavioral state-dependent: they remain silent during simulated head movement in forward locomotion, but they provide identical and constant output when simulating reversal.

The combined neuronal outputs activate the muscle and drive the head bending:

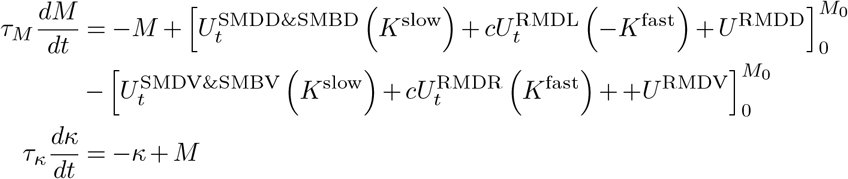

Positive and negative values of *M* represent dorsal and ventral muscle activation, respectively. A clipping function 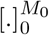 limits the muscle activation level between 0 and *M*_0_. The RMDL senses ventral bending and drives dorsal bending, contributing 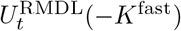. Conversely, RMDR neurons contribute 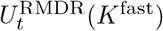 for ventral bending. The coefficient *c* scales the contribution of the RMD group..

We determined the head’s mechanical time constant *τ*_*κ*_ through relaxation experiments. Following optogenetic activation of unilateral head muscles, we analyzed the decrease in the head curvature after the termination of head muscle activation. By fitting exponential functions to these relaxation curves, we found that the head returns to its resting position with a characteristic time scale of 0.4 seconds. This rapid relaxation is notably faster than the body’s mechanical time constant (48), likely due to the head’s lower mechanical load. The head’s quick response dynamics prove essential for generating the rapid head casts captured in our model. The time constant for muscle activation *τ*_*M*_ is 0.1 seconds following previous works(50, 56).

We examined how well our model’s surrogate posterior curvature aligned with the actual posterior curvature of worms. To do this, we compared two time lags: (1) the lag between head and neck curvature in N2 worms, and (2) the lag between head and surrogate posterior curvature in our model. We determined these lags by identifying the time point at which the cross-correlation between the head curvature’s slow mode and posterior curvature signal reached its maximum.

Simulated RMD ablation was implemented by removing the contributions from 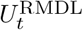 and 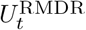 by setting *c* = 0. The simulated neck inhibition was implemented by scaling the slow feedback *K*_slow_ by setting *a*_*s*_ ← 0.6*a*_*s*_.

### Neuro-mechanical Model

To investigate the impact of the temporal pattern of head bending on the curvature amplitude of the rest of the body, we employed a bio-realistic mechanical model from (50) to simulate forward locomotion. We proposed a ventral nerve cord (VNC) circuit featuring bistable motor neuron pairs and anterior proprioceptive coupling to drive this mechanical model.

The mechanical model (50) comprises 48 connected mechanical units, with the body wall muscles modeled as elastic rods capable of active contraction. The model is controlled by 12 pairs of dorsal and ventral motor neurons, with each pair governing 4 mechanical units.

The dynamics of each pair of body motor neurons is governed by two main mechanisms:

1. Dorsal contraction in the anterior units activates the dorsal motor neuron, and ventral contraction activates the ventral motor neuron.
2. Reciprocal inhibition exists between the dorsal and ventral motor neuron partners.

We use directional proprioceptive coupling (32) to induce an activation of motor neurons through anterior bending curvature changes(Fig. S5A). The mutually inhibiting neuron pairs operate within a bistable regime, modulated by the curvature in the anterior section of the body (Fig. S5B).

Let *S*_*t*_ = 1 represent the dorsal domination, and *S*_*t*_ = −1 represent the ventral domination. The proprioceptive signal *P*, which is equal to the mean curvature of anterior 8 units (48 units in total), switches the motor neurons pair between dorsal domination and ventral domination:

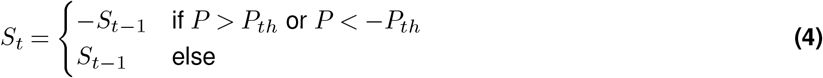

Here, *P* is positive for anterior dorsal bending and negative for ventral bending. When *S* = 1, the dorsal motor neuron dominates (Fig. S5B upper and lower branches for Vd and Vv, respectively):

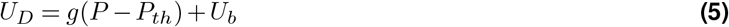

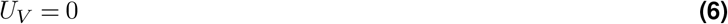

Conversely, when *S* = −1, the ventral motor neuron is activated (Fig. S5B lower and upper branches for Vd and Vv, respectively):

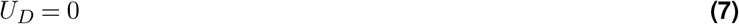

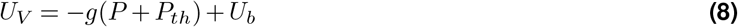

Here *U*_*D*_ and *U*_*V*_ denote the activation levels of the dorsal and ventral motor neurons, respectively. *U*_*b*_ is the baseline activation and *g* modulates how the activation level changes with proprioception. We set *g >* 0 so that deeper anterior bending leads to higher motor neuron activity and thus deeper posterior bending. This activation function helps to propagate the change in the head bending amplitude along the body.

Finally, the activation of the foremost motor neuron pair, which drives head bending, is controlled by an external input *u*_*t*_, which alternates between 1 and -1 i.e. between dorsal activation and ventral activation:

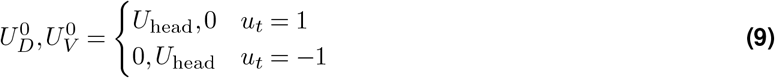

The head input is simplified to a binary signal to capture rapid head casts, which are more likely due to abrupt changes in head motor neuron output rather than gradual gain modulation (38, 40).

Using our neuro-mechanical model, we can simulate forward locomotion by applying periodic input to the head motor neurons. Our goal is to identify the input pattern that maximizes kinematic efficiency, defined as the ratio of the average locomotion speed to the undulation frequency. This efficiency measure represents the distance the worm covers in one undulation period.

To explore possible input patterns, we discretized one period of the head input signal into 20 intervals, resulting in a binary sequence of length 20. To further constrain the search space, we imposed dorsal-ventral symmetry around the midpoint and limited the number of transitions between dorsal and ventral activation to no more than two in half a period. These constraints reduced the number of possible input signal profiles to 72. Among these profiles, we identified the one that yielded the highest kinematic efficiency.

### Optimal angle of attack for maximum locomotion speed

We describe the traveling body wave along the moving direction with 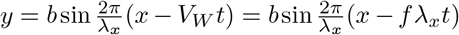, where *b* is the body wave amplitude, *λ*_*x*_ is the effective wavelength along the moving direction, *V*_*W*_ = *fλ*_*x*_ is the wave speed, and *f* is the undulation frequency. For simplicity, the wave number is 1 in this derivation.

It’s crucial to distinguish between the effective wavelength and the wavelength in the body length coordinate used in previous works (47, 48). While these wavelengths are similar for small amplitudes, they diverge as the amplitude increases. This divergence occurs because the effective wavelength decreases due to the shrinkage along the anterior-posterior axis in a more curled posture. The sine wave amplitude *b* and effective wavelength *λ*_*x*_ are related by a constant worm length:

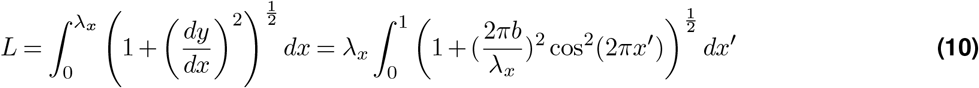

Follow Gray and Hankcock (47), the resistive force on an element *ds* along the moving axis is given by:

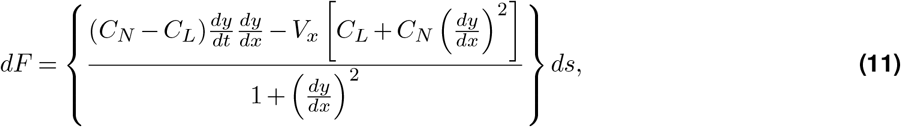

For large amplitude, we have 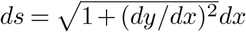 thus:

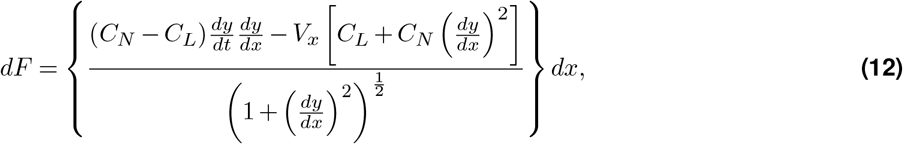

Total force over a complete wave is:

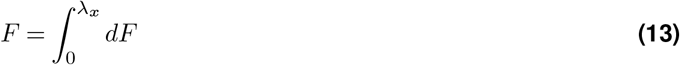

We introduce integrals:

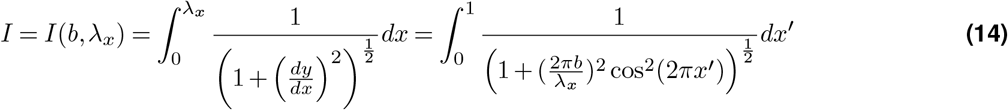

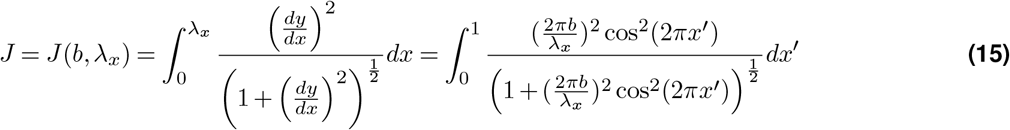

Using the zero net force condition:

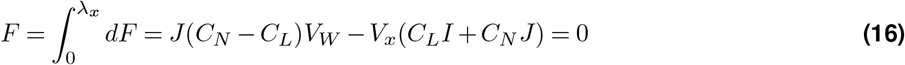

The locomotion speed is derived as:

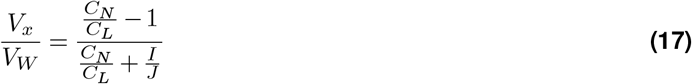

We then can numerically calculate the moving speed of the worm at an arbitrarily varying amplitude *b*, with a fixed frequency and with the wave number equal to 1:

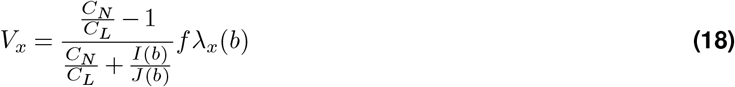

To further simplify the equation, we apply a small amplitude approximation to the integrals:

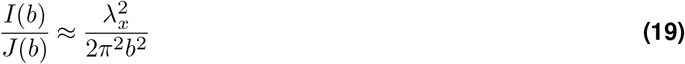

The body angle *α* along the moving direction follows 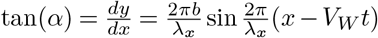. The angle of attack *A*_*a*_ is defined as the maximum angle, which follows

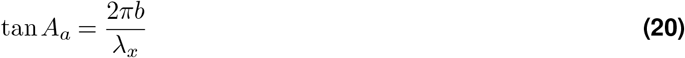

Combining these, we arrive at:

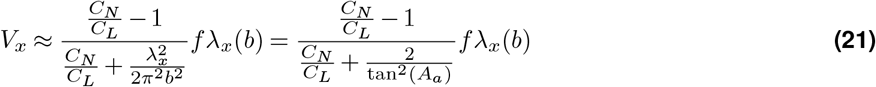

Our numerical analysis suggested that Eq. (21), which incorporates the small amplitude approximation in the integrals and wavelength compensation (accounting for finite amplitude), closely approximates Eq. (18). We defined the kinematic efficiency as:

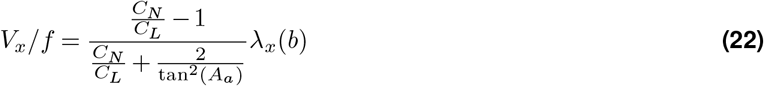

that equals the distance covered by the worm in one period of undulation.

## Supplementary figures (S1)

**Figure S1.**
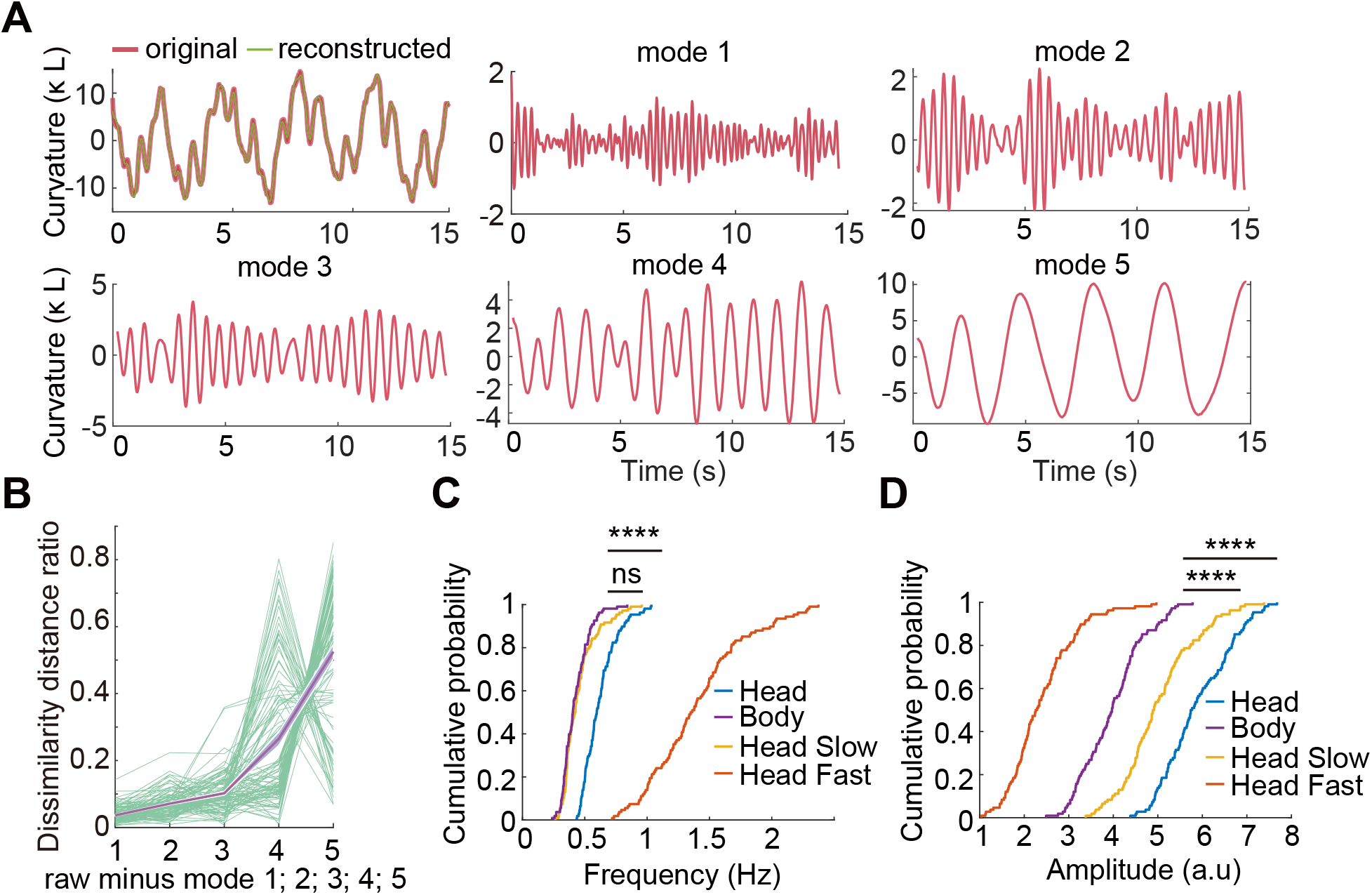
Fast and slow head movement dynamics. **A**. Representative time series of 5 modes from Variational Mode Decomposition (VMD) of a head curvature signal. **B**. Contribution of each decomposed mode to head motion signal, higher dissimilarity represents higher contribution. **C, D**. Cumulative probability distribution of bending frequency and amplitude of head and body movements, as well as slow and fast dynamic mode of head curvature. K-S test. n.s. p > 0.05, ****p < 0.0001,

**Figure S2.**
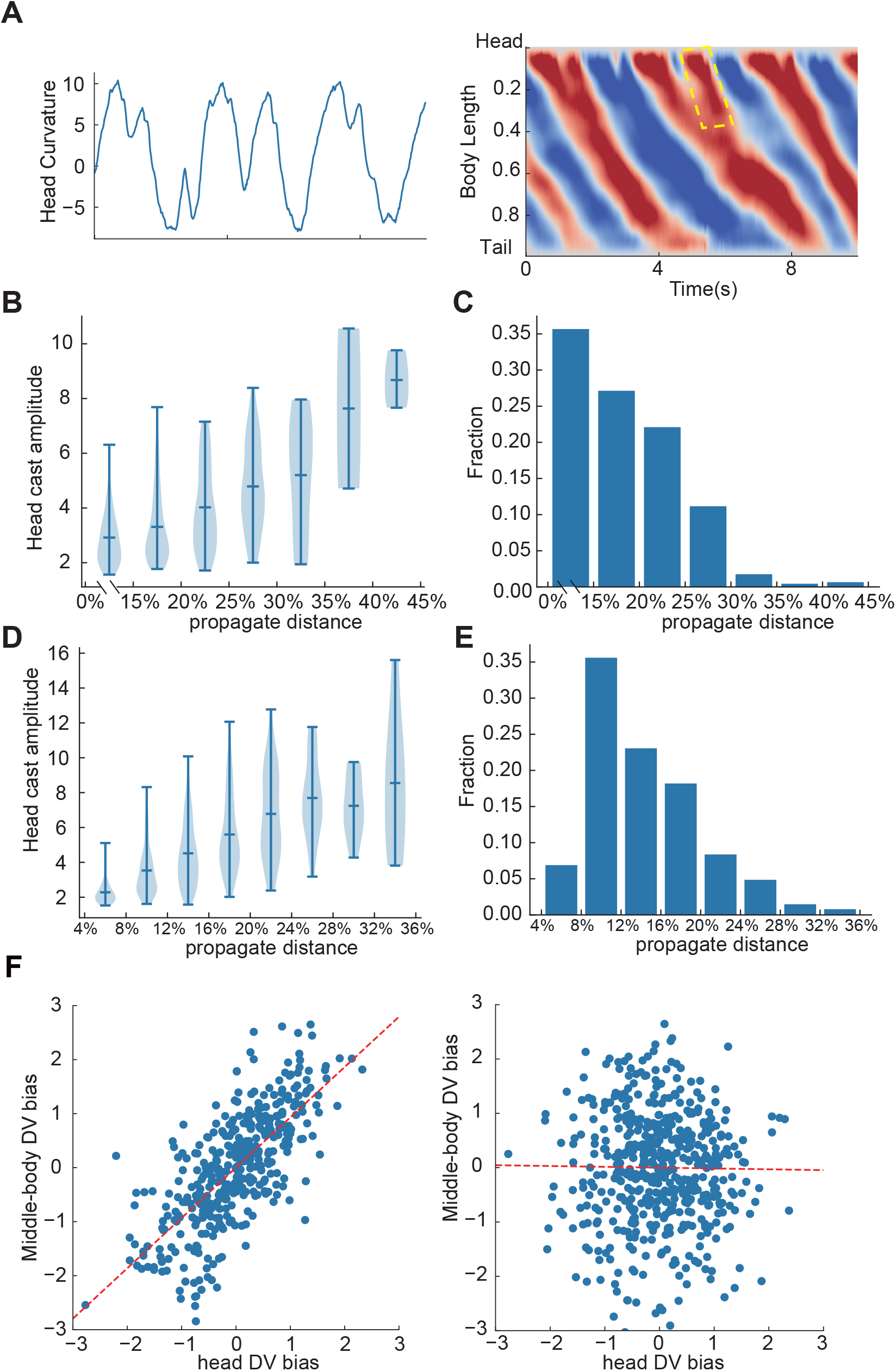
Head cast propagation. **A**. Kymograph demonstrating the propagation of head casts to the middle body (highlighted by yellow dashed box). **B**. Correlation between head cast amplitude and propagation distance. Larger head casts exhibit greater propagation distances. **C**. Distribution of head cast propagation distances. **D-E** Similar to **B** and **C**, the head casts were identified using curvature from the anterior 4% - 8% body length instead of 0% - 15%. **F**. Analysis of dorsal-ventral (D-V) bias correlation between head and middle-body curvatureLeft: Strong correlation between D-V bias in head curvature and middle-body curvature when head undulation propagates to the middle body (r=0.73). Right: No correlation was observed when comparing head D-V bias to middle-body curvature from preceding propagation (control, r=-0.01).

**Figure S3.**
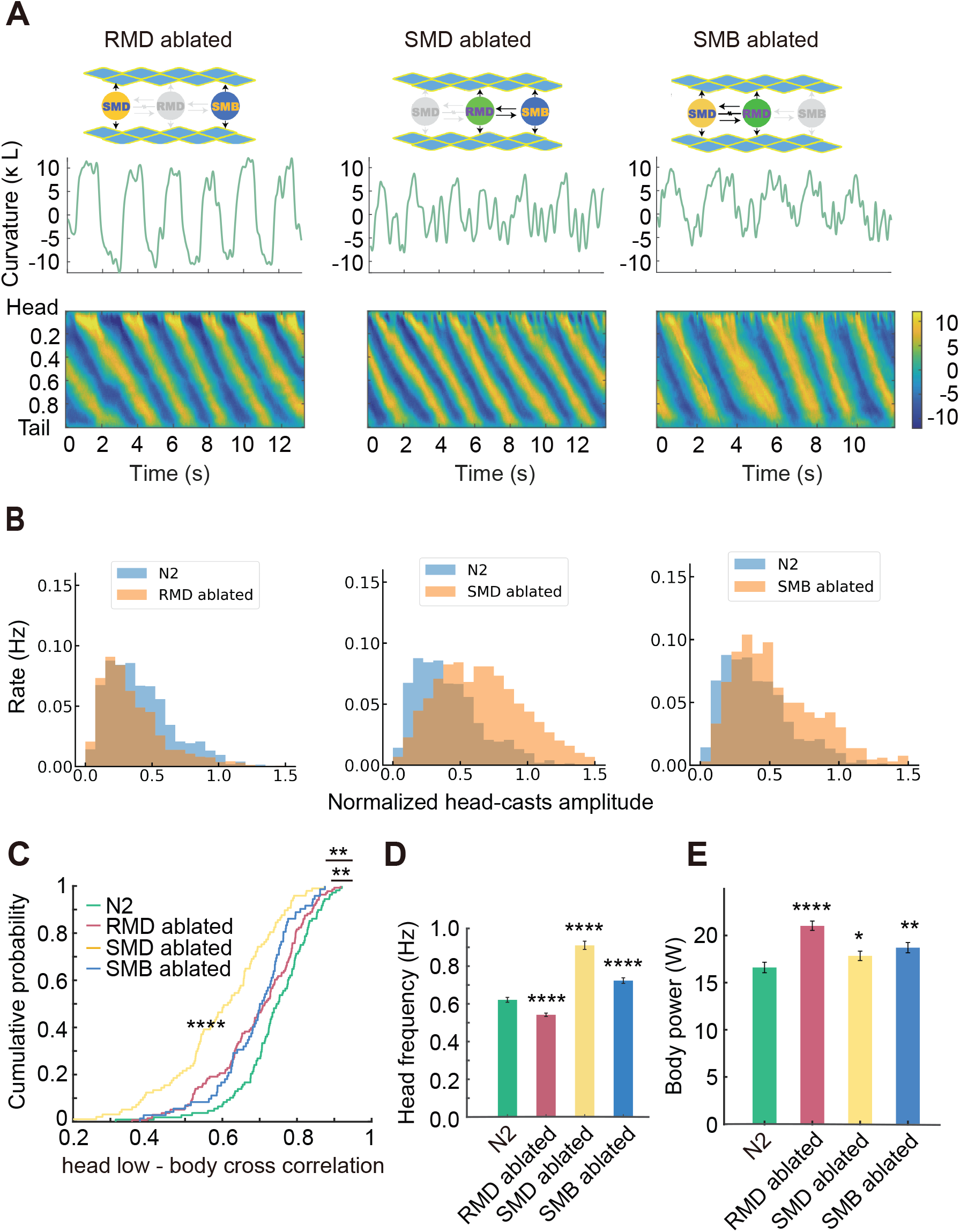
Head motor neuron ablation. **A**. Ablation schemes (top), illustrative head curvature time series (middle), and whole body curvature kymogram (bottom) for RMD, SMD, and SMB ablated animals. **B**. Occurring rate of head casts at different relative amplitude for RMD, SMD, and SMB ablated worms. **C**. Cumulative probability distribution of cross correlation coefficient between middle body curvature and the slow mode of head curvature. KS test. **D**. Head undulation frequency changes for RMD, SMD, and SMB ablated animals. **E**. Body power changes for RMD, SMD, and SMB ablated animals. Mann-Whitney U test. *p < 0.05**p < 0.01, ****p < 0.0001,

**Figure S4.**
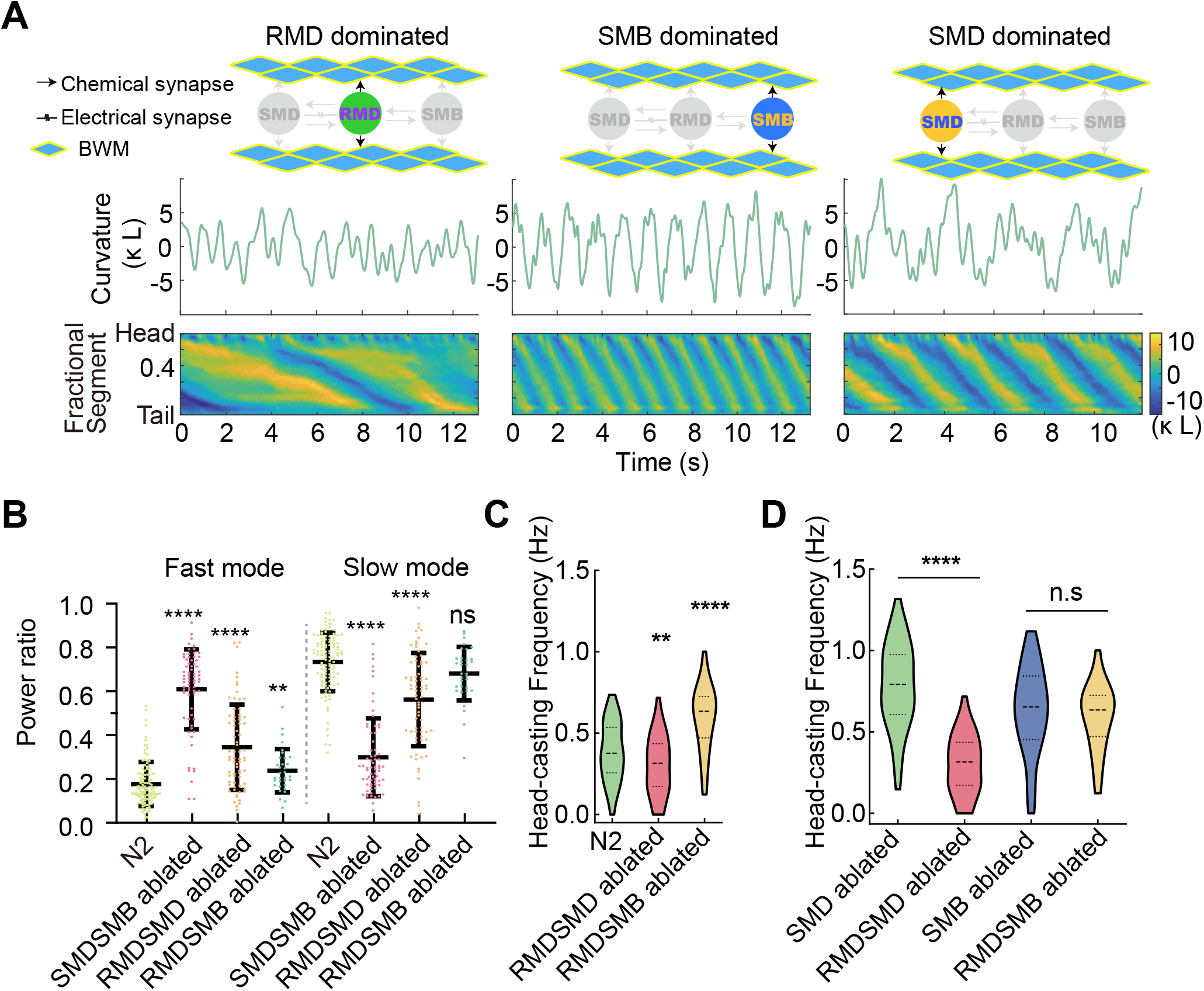
Head motor neuron co-ablation. **A**. Ablation schemes (top), example curvature time series (middle), and whole body curvature kymogram (bottom) for SMD&SMB, RMD&SMD, and RMD&SMB co-ablated animals. **B**. Power ratio of fast mode and slow mode to total signal power for co-ablated animals. **C-D**. head-casts frequency changes for RMD&SMD, and RMD&SMB co-ablated animals. n.s. p > 0.05, **p < 0.01, ****p < 0.0001, Mann-Whitney U test.

**Figure S5.**
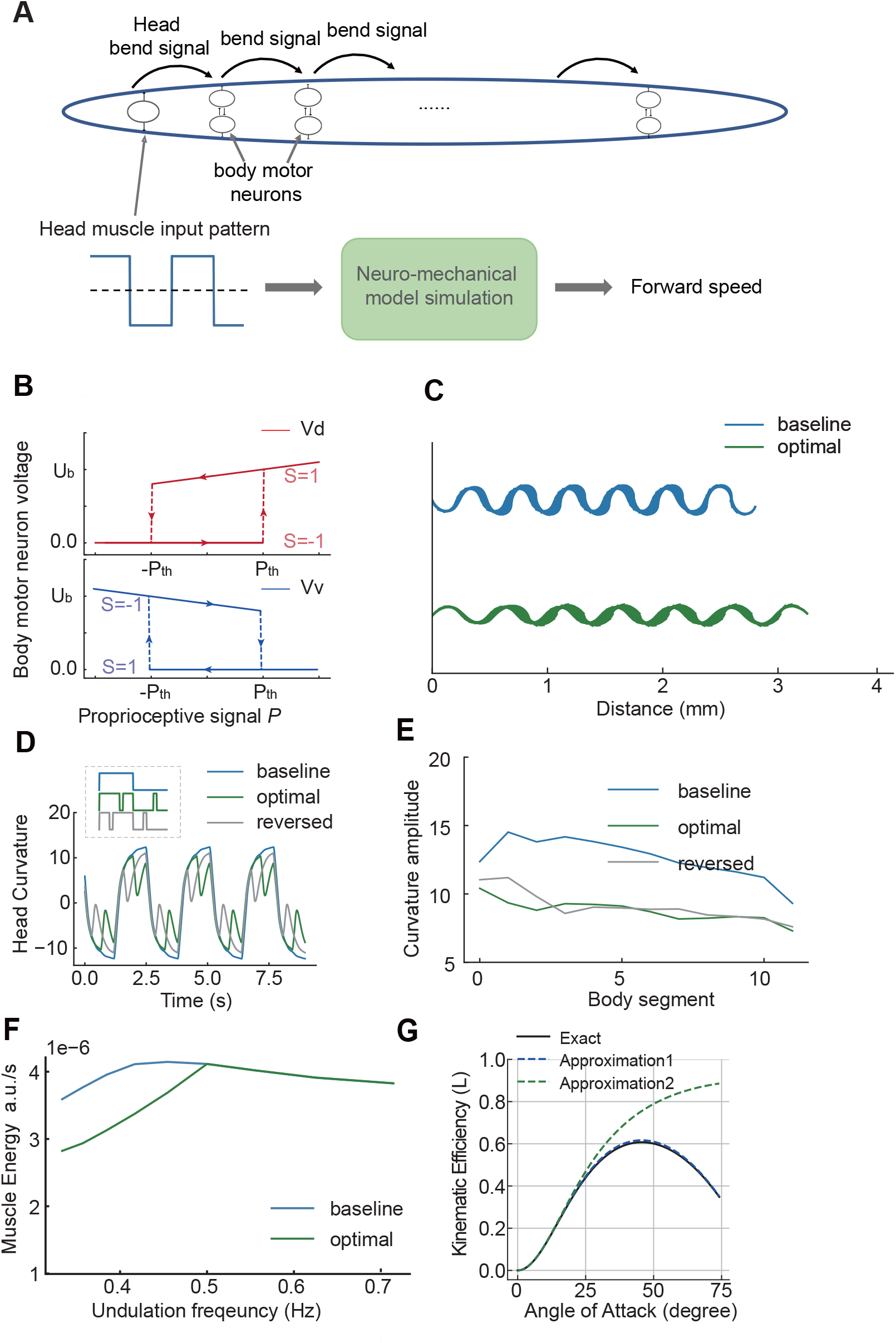
Kinematic efficiency. **A**. Head muscle input pattern drives head bending. Body motor neuron pairs nonlinearly transform anterior curvature to local muscle activation, propagating the bending. The mechanical model (50) receives muscle activation from the motor circuit to generate forward locomotion. We measure locomotion speed and muscle output power. **B**. Two hysteresis curves model the mutually inhibiting motor neuron pairs. Proprioceptive signals switch their state between two branches (S=1 and S=-1). **C**. Crawling trajectories from simulation with baseline and optimized head input in Fig. 6E. **D-E**. Reversing the optimized head input pattern (gray line) increases anterior curvature, resulting in sub-optimal kinematic efficiency. **F**. Estimated Muscle Energy Expenditure: Comparison of baseline and optimized head input across various undulation frequencies. **G**. Theoretical calculation of kinematic efficiency. Two theoretical approximations of kinematic efficiency using small amplitude assumptions. Approximation 1 includes wavelength correction, while Approximation 2 does not.

**Figure S6.**
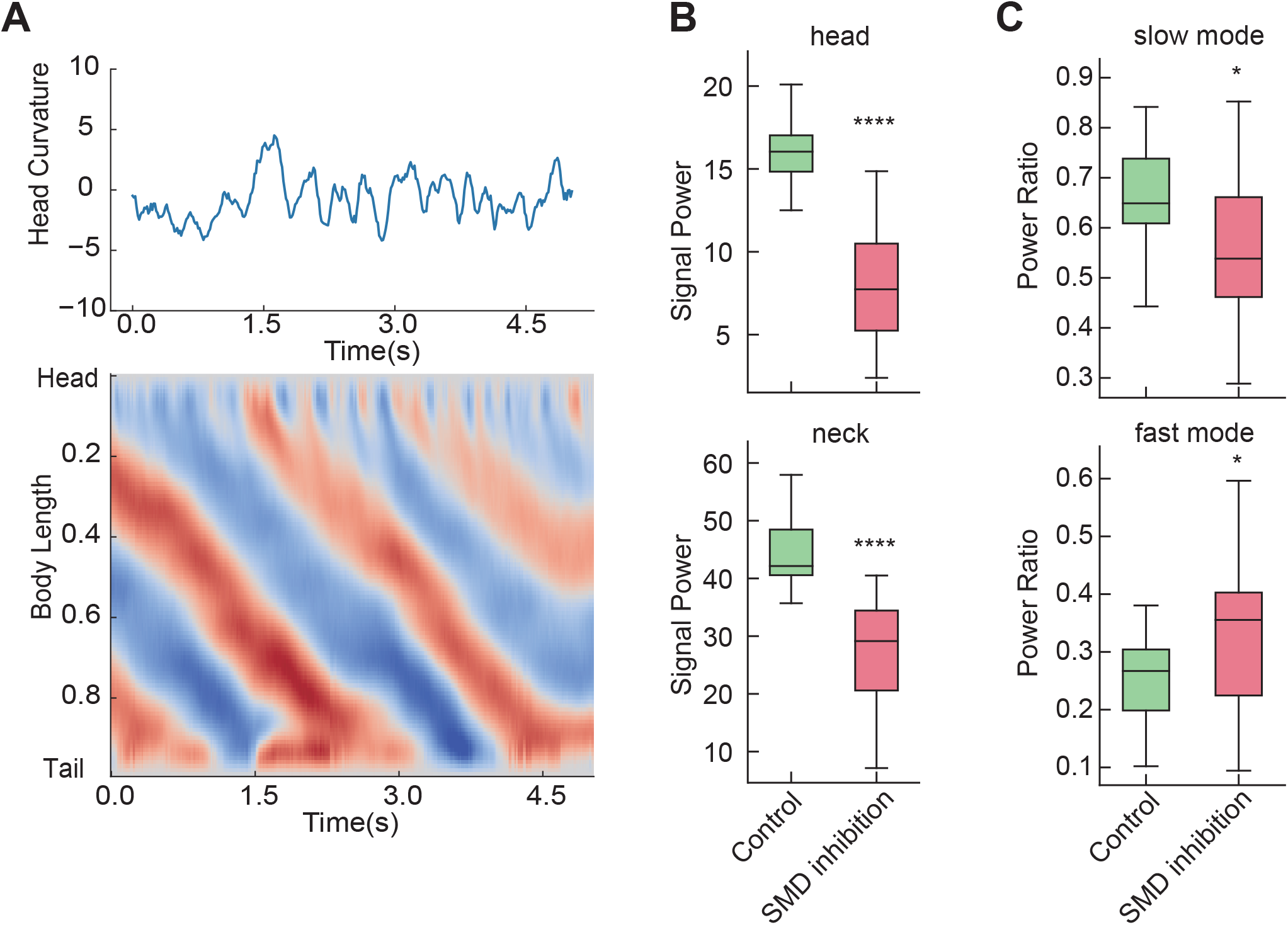
SMD inhibition. **A**. Representative head curvature time series (top) and whole body curvature kymogram (bottom) during SMD inhibition. **B**. SMD inhibition resulted in a significant decrease in head and neck curvature, quantified by curvature signal power. **C**. SMD inhibition resulted in an increased power ratio in fast mode and a decreased ratio in slow mode. *p < 0.05, ****p < 0.0001, Wilcoxon signed-rank test.

**Figure S7.**
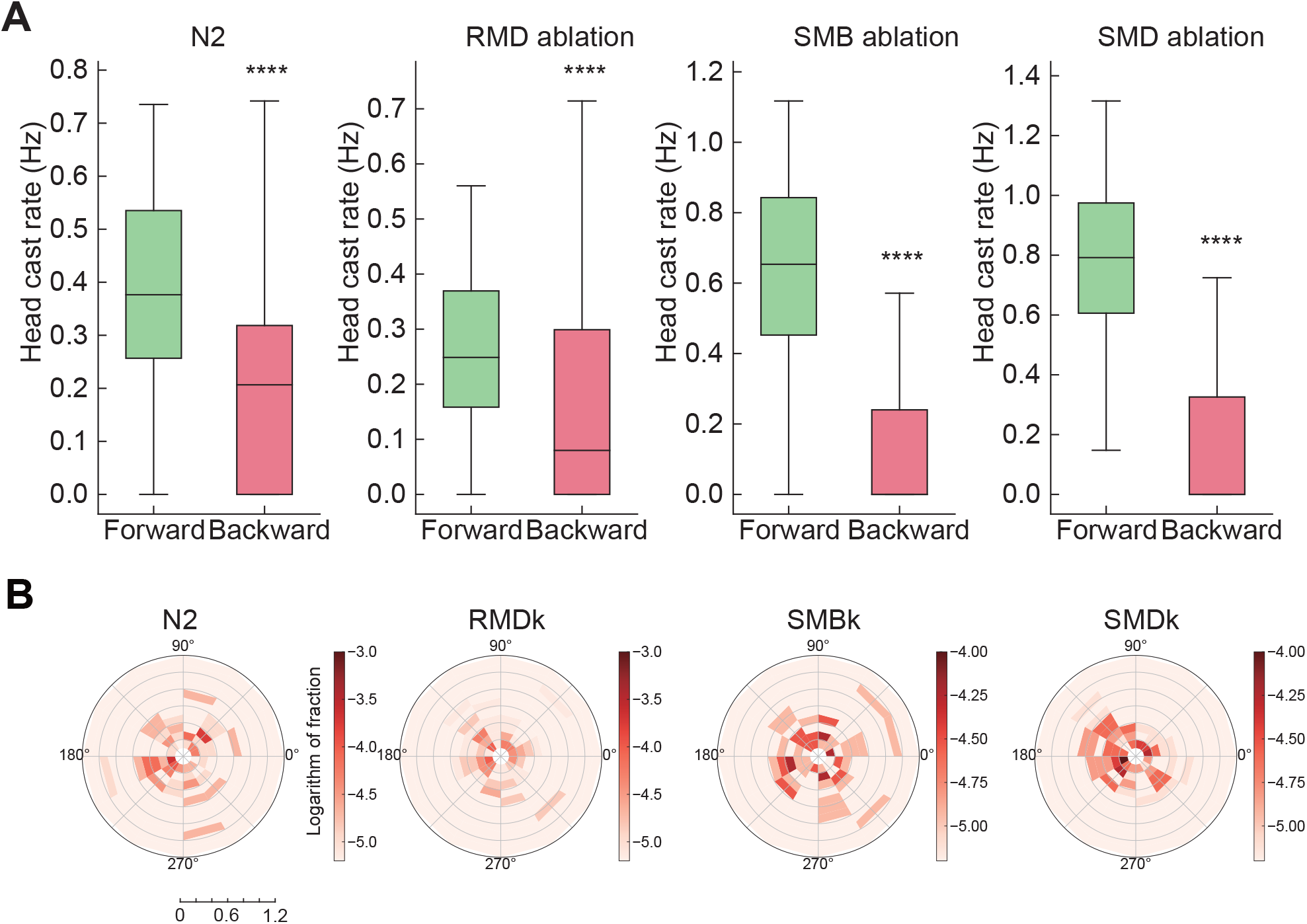
Head-cast statistics during reversal. **A**. Reduced head casts rate during reversals in N2 and head motor neuron ablated animals. **B**. Phase-amplitude distribution of head casts during reversals in N2 and mutants. ****p < 0.0001, Mann-Whitney U test.

**Figure S8.**
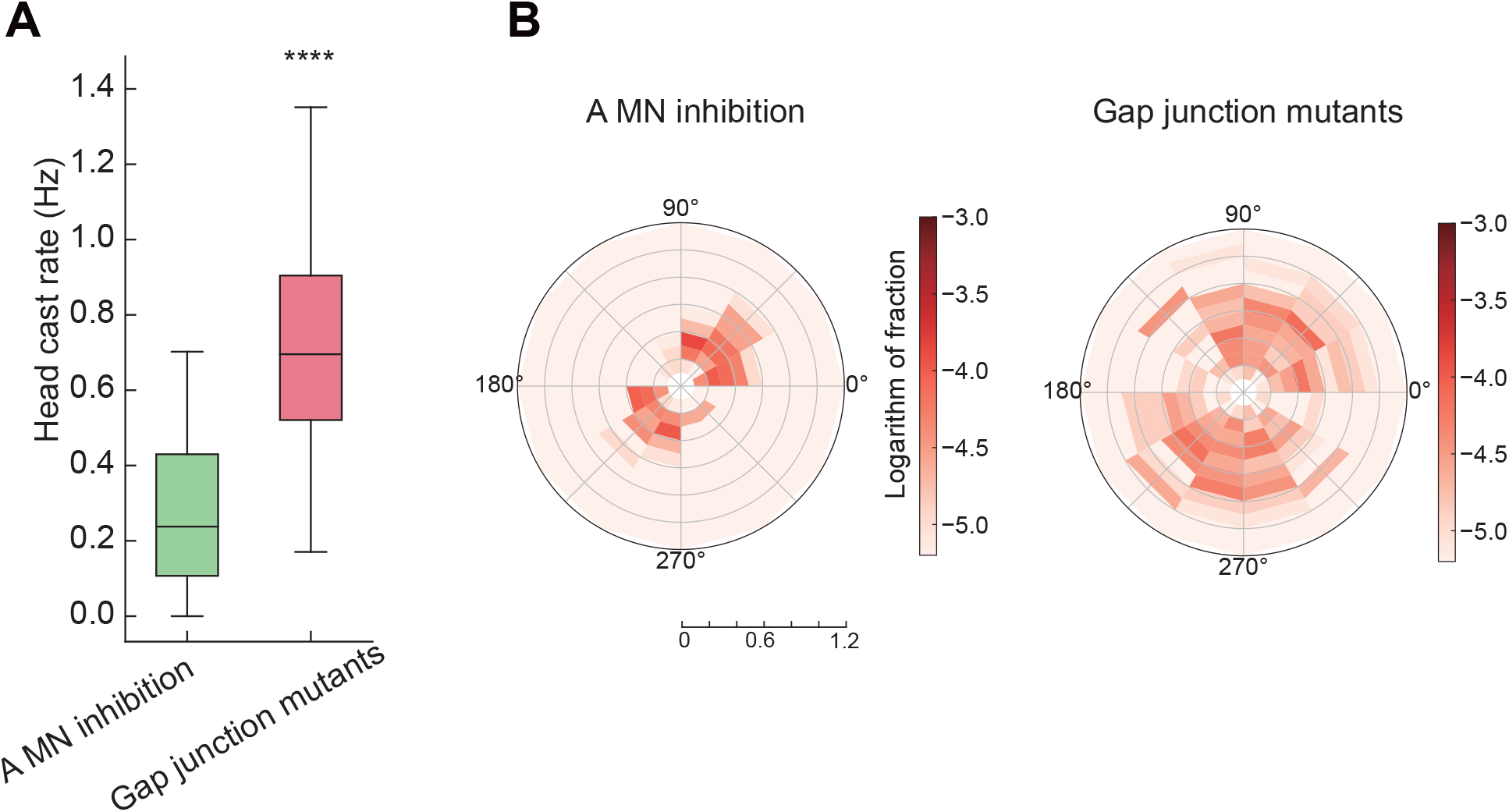
Phase coordination are disrupted in gap junction mutants. Gap-junction deficient mutants (*unc-7* ;*unc-9*) showed increased head casts (**A**) and disrupted phase specificity (**B**). To promote forward movements in these animals, we silenced A-type motor neurons (P*unc-4::twk-18(gf)*) to suppress backward locomotion (31). ****p < 0.0001, Mann-Whitney U test.

## Supplementary tables (S2) Information of plasmids and strains

**Table S1.**
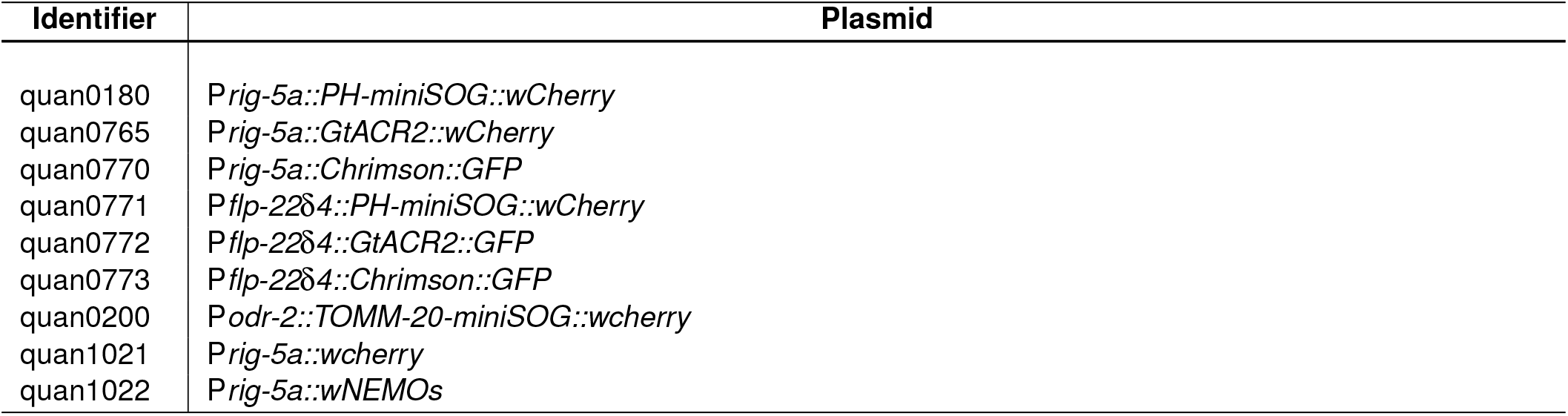
Plasmids used in this work.

**Table S2.**
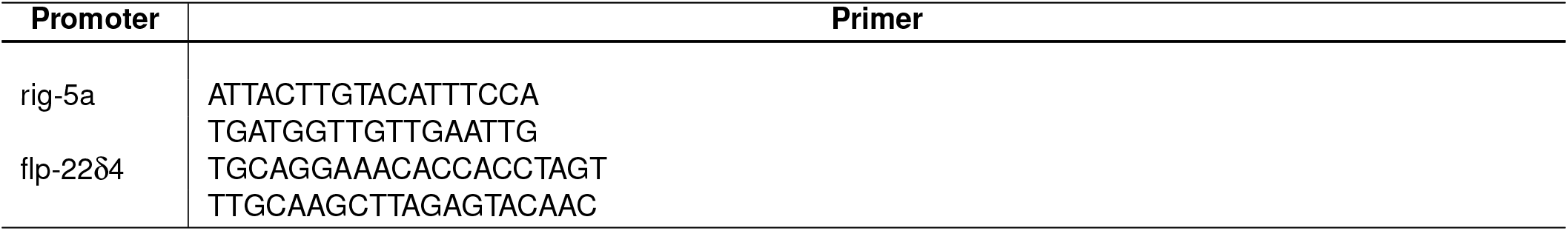
Primers used in this work.

**Table S3.**
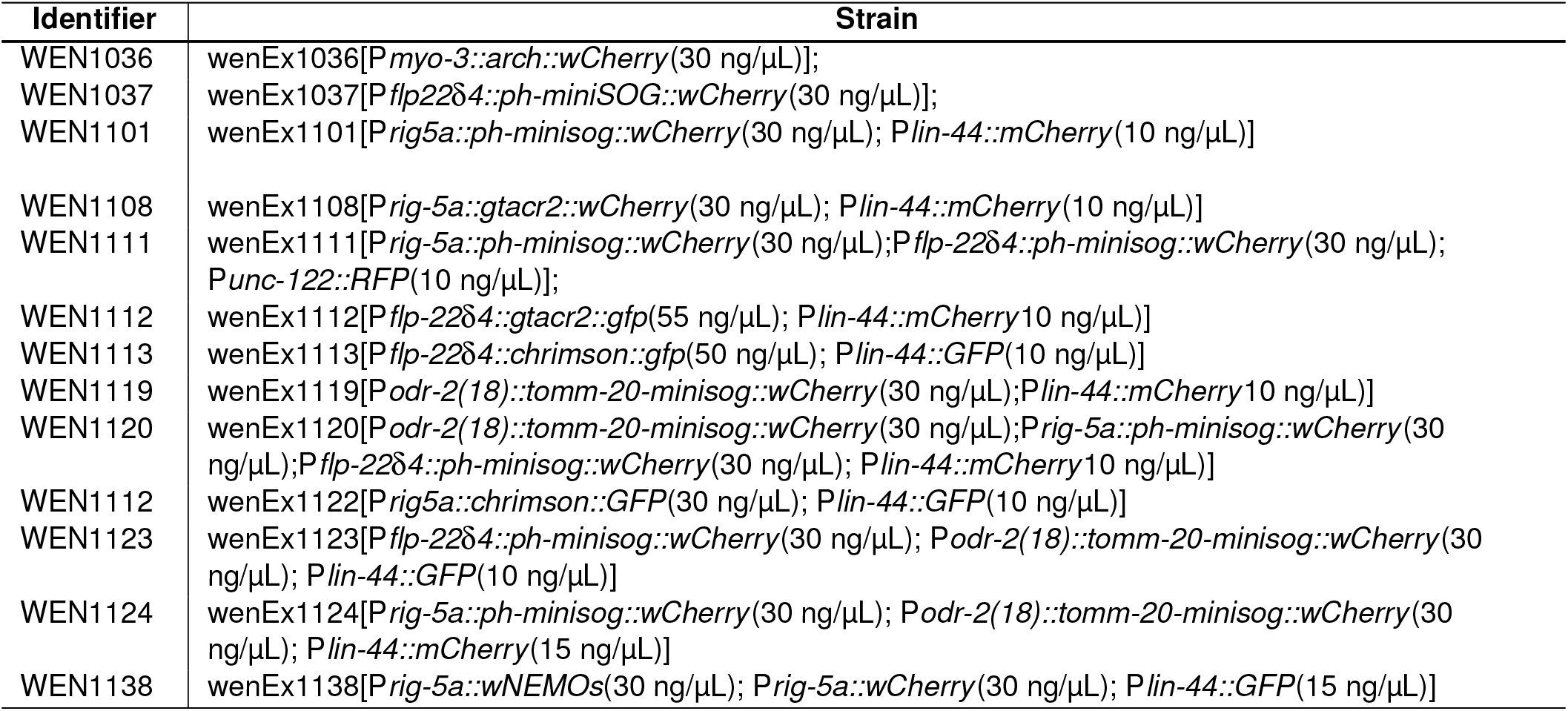
Strains used in this work.

